# Striatal dopamine dynamics across temporal scales in brain-specific *Atp2a2* heterozygous knockout mice during reward conditioning

**DOI:** 10.64898/2026.09.01.748702

**Authors:** Mizuho Niido, Takamasa Yoshida, Noriko Isoo, Masakazu Mimaki, Kazuo Nakajima, Toshihiro Hayashi

## Abstract

Dysregulated dopamine signaling is implicated in schizophrenia and mood disorders, yet how disease-associated genes alter dopamine dynamics during behavior remains poorly understood. *ATP2A2* encodes sarco/endoplasmic reticulum Ca^2+^-ATPase 2 (SERCA2), a regulator of intracellular Ca^2+^ homeostasis, and has been associated with psychiatric vulnerability. Brain-specific *Atp2a2* heterozygous conditional knockout (het-cKO) mice exhibit behavioral abnormalities and elevated extracellular dopamine in the nucleus accumbens (NAc) with microdialysis, but it lacks the temporal resolution to resolve rapid, context-dependent dopamine dynamics. Here, we used dual-site fiber photometry with the genetically encoded dopamine sensor dLight1.2 to monitor dopamine signals simultaneously in the NAc and tail of the striatum (TS) during classical reward conditioning. Control and het-cKO mice acquired task-related licking comparably. Nevertheless, het-cKO mice showed enhanced NAc reward-evoked dopamine responses in the pooled analysis, with indications of more frequent and longer-lasting spontaneous transients. Responses to unexpected reward omission were larger in het-cKO mice in both regions during the early phase, but they were not detected after further training. Notably, late-phase suppression of reward-predictive NAc dopamine responses by a preceding aversive outcome persisted in het-cKO mice, although evidence for genotype dependence was inconclusive. Regional dopamine transporter (DAT) protein expression showed no clear genotype differences. These findings identify context-dependent alterations in dopamine signaling across multiple temporal scales and striatal regions in brain-specific *Atp2a2* haploinsufficiency, highlighting intracellular Ca^2+^ regulation and dopamine release as priorities for direct investigation. These results may help clarify the biological links between *ATP2A2* haploinsufficiency and neuropsychiatric dysfunction.

## Introduction

Dysregulation of dopamine signaling is central to several psychiatric disorders, including schizophrenia and mood disorders [1, 2]. Whereas early dopamine hypotheses emphasized postsynaptic receptor abnormalities, current evidence points more strongly to presynaptic dysfunction involving dopamine synthesis, release, and temporal dynamics [3, 4]. Large-scale genetic studies have also identified numerous psychiatric-risk loci [5], but how these genes perturb dopamine signaling during behavior remains unclear. Genetic vulnerability may alter not only overall dopamine tone but also when, where, and in what behavioral context dopamine signals are generated.

*ATP2A2* is a plausible molecular link between psychiatric risk and dopamine dysfunction. As a sarco/endoplasmic reticulum Ca^2+^ pump, ATP2A2 transports cytosolic Ca^2+^ into the endoplasmic reticulum (ER) and thereby contributes to intracellular Ca^2+^ homeostasis. Heterozygous mutations including loss-of-function variants in *ATP2A2* cause Darier disease, an inherited skin disorder associated with increased prevalence of schizophrenia, bipolar disorder, major depression, and other neuropsychiatric symptoms [6–9]. Brain-specific *Atp2a2* heterozygous conditional knockout (het-cKO) mice were previously shown to exhibit delayed neuronal Ca^2+^ clearance, altered behavioral responses to novel environments, impaired fear memory, and elevated extracellular dopamine in the nucleus accumbens (NAc) [10]. These observations suggest that partial loss of ATP2A2 function can influence both neuronal Ca^2+^ regulation and mesolimbic dopamine signaling.

The previous dopamine measurements in het-cKO mice were obtained by microdialysis [10]. Although this approach established a hyperdopaminergic state in the NAc, its minute-scale temporal resolution could not resolve rapid fluctuations during individual behavioral events. Dopamine signals span several temporal domains that convey distinct information. Subsecond transients encode reward prediction errors and contribute to associative learning [11]. Fluctuations over seconds reflect ongoing network activity and are shaped by both release and extracellular clearance [12, 13]. Across successive trials, dopamine responses can also integrate recent outcomes over distinct temporal horizons, allowing current cues to be interpreted in light of recent experience [14]. ATP2A2 deficiency could therefore alter dopamine signaling at one or several of these levels, even when overall task performance is preserved.

Dopamine dynamics also differ across striatal circuits. NAc-projecting dopamine neurons are linked to reward prediction, motivational value, and approach behavior, whereas TS dopamine is prominently recruited by novel and threatening stimuli [15–17] and can convey prediction-error signals during associative fear learning [18]. Comparing these regions therefore tests whether *Atp2a2* haploinsufficiency broadly disrupts dopamine signaling or selectively affects temporal components across circuits.

Here, we used dual-site dLight1.2 fiber photometry to monitor NAc and TS dopamine signals simultaneously in control and *Atp2a2* het-cKO mice during classical reward conditioning and pharmacological validation. We quantified task-evoked responses, spontaneous transients, spectral organization, and trial-history-dependent modulation to determine how *Atp2a2* haploinsufficiency reshapes dopamine signaling across temporal scales and striatal circuits.

## Materials and Methods

Complete methodological details are provided in the Supplementary Materials and Methods.

### Animals

Brain-specific *Atp2a2* het-cKO mice were generated as described previously [10]. *Atp2a2*^flox/+^; *NesCre*^tg/+^ mice constituted the het-cKO group and *Atp2a2*^+/+^; *NesCre*^tg/+^ littermates served as controls. Genotyping primers are listed in Supplementary Table S1. Fiber-photometry experiments included 10 mice per genotype (control, 4 males/6 females; het-cKO, 5 males/5 females), aged 16.6–50.4 weeks at the first Early-phase recording (Supplementary Table S2). All procedures were approved by the Animal Care and Use Committee of Teikyo University School of Medicine (No. 24-018) and the Teikyo University Genetically Modified Organism Experimental Safety Committee (No. 2404382A1) and were conducted in accordance with the approved guidelines.

### Surgery and fiber photometry

AAV5-hSyn-dLight1.2 (1.0 × 10^13^ GC/mL; Addgene #111068-AAV5) was injected into the left NAc (AP +1.2, ML +1.0, DV −4.5 mm) and right TS (AP −1.0, ML −3.25, DV −2.5 mm) [17]. One microliter was delivered at each site, followed by implantation of a 400-µm-core, 0.39-NA optical fiber. Recordings began ≥2 weeks later using simultaneous dual-site interleaved 405/470-nm excitation at 40 Hz per wavelength with a camera-based system (Fig. 1c) [19, 20].

**Figure 1.**
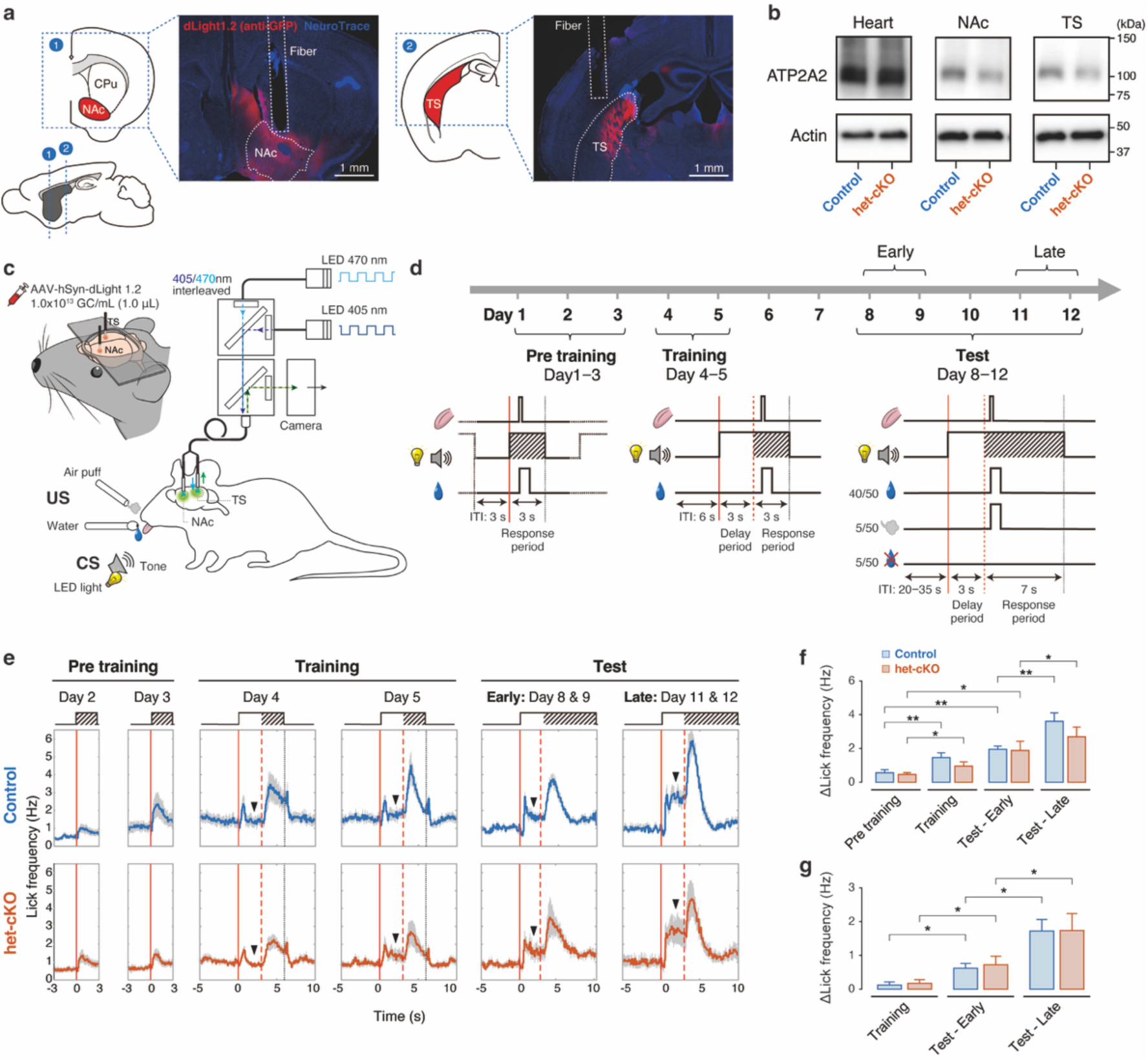
*Atp2a2* het-cKO mice, fiber-photometry recording, and behavioral paradigm. (a) Schematics and representative histological images showing AAV-hSyn-dLight1.2 expression and optical fiber placements in the left nucleus accumbens (NAc; site 1) and right tail of the striatum (TS; site 2). dLight1.2 was visualized by GFP immunostaining (red) with NeuroTrace counterstaining (blue). Scale bars, 1 mm. CPu, caudate–putamen. (b) Representative western blots of ATP2A2 and β-actin in the NAc, TS, and heart from one control and one het-cKO mouse. Quantitative reduction of brain ATP2A2 expression in this het-cKO line was established previously [10]. (c) Fiber-photometry and behavioral apparatus. CS, conditioned stimulus; US, unconditioned stimulus. (d) Task schedule: Pre-training (days 1–3), Training (days 4–5), and Test (days 8–12). Days 8 and 9 were defined as the Early phase, and days 11 and 12 as the Late phase. Day 10 was an intervening Test session and was not included in the phase-based analyses. Test sessions comprised 40 Reward, 5 Air-puff, and 5 Omission trials. (e) Lick-frequency peri-stimulus time histograms (50-ms bins, no smoothing; mean ± SEM). Solid and dashed red lines indicate CS onset and response-period onset, respectively; arrowheads indicate the mean lick-frequency levels during the delay period. (f) Response-period Δlick frequency, defined as the mean during the first 2 s of the response period minus the mean during the −2- to 0-s baseline. (g) Delay-period Δlick frequency during the common 1–3-s window relative to the same baseline. For (f, g), values were averaged across two sessions within each phase for each mouse (*N* = 10 mice per genotype); bars show means with upper SEM error bars across mice. Genotype × Phase repeated-measures ANOVA with Holm-adjusted comparisons. \**p* < 0.05; \*\**p* < 0.01 (Holm-adjusted).

### Behavioral task

Mice underwent head-fixed classical conditioning adapted from appetitive and aversive paradigms (Fig. 1c) [21, 22]. The CS combined a 10-kHz amplitude-modulated tone and flashing blue LED. Test outcomes were 3–5 µL of 0.1% saccharin, a 0.5-s air puff, or omission (80%, 10%, and 10%). The schedule comprised Pre-training (days 1–3), Training (days 4–5), and Test (days 8–12); days 8–9 were Early, days 11–12 Late, and day 10 was an intervening Test session excluded from phase analyses (Fig. 1d). Licks were detected by infrared-beam interruption and baseline-subtracted relative to −2 to 0 s. Detailed task parameters are provided in Supplementary Table S3.

### Pharmacological validation

Pharmacological validation began on day 12, whose Test session served both as Late-phase session 2 and the Before-drug baseline for the first GBR12909 administration. GBR12909 (20 mg/kg, i.p.) inhibited DAT [23], whereas SCH23390 (0.25 mg/kg, i.p.) blocked D1 receptors [12]. Each experiment compared a 50-trial Before session with a second Test session 60 min after injection. GBR12909 was administered on days 12 and 17 and SCH23390 on days 15 and 19; separate drug-free Test sessions were conducted on days 16 and 18 (Fig. 2c). The complete experimental-day assignments and analytical roles of each session are summarized in Supplementary Table S4.

**Figure 2.**
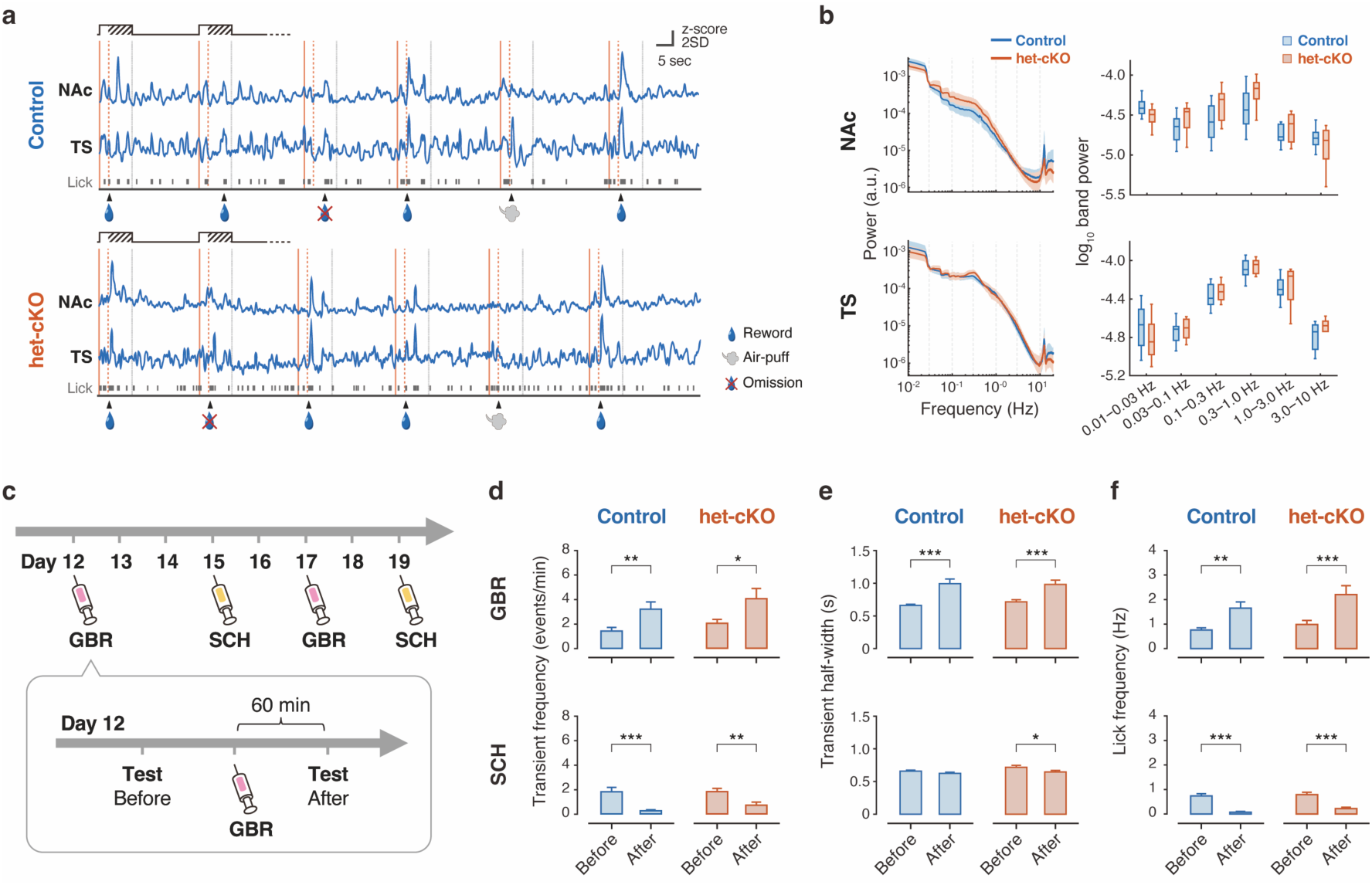
Striatal dopamine signals and pharmacological validation. (a) Representative z-scored dopamine traces recorded simultaneously from the NAc and TS during six consecutive trials. Lick events and trial outcomes are shown below the traces; arrowheads indicate the first lick during the response period. (b) Power spectra and band power in six frequency bands. Spectral traces show means and 95% confidence intervals. Box plots show the median, interquartile range, minimum-to-maximum whiskers, and individual mice (*N* = 10 mice per genotype). Genotype comparisons were performed using mouse-level averages. (c) Pharmacological schedule. Before- and After-drug recordings were separated by 60 min. The day 12 Before-drug Test session was the same recording used as the second Late-phase session. GBR, GBR12909; SCH, SCH23390. (d–f) Effects of GBR12909 (top) and SCH23390 (bottom) on NAc ITI transient frequency (d), transient half-width (e), and ITI lick frequency (f). Complete matched Before–After session pairs were analyzed using session-level linear mixed-effects models with MouseID and PairID as random effects. Bars show session means with upper SEM error bars. Maximum sample sizes were *n* = 17 matched pairs from *N* = 7 mice (control, GBR12909), *n* = 16 from *N* = 7 mice (control, SCH23390), *n* = 16 from *N* = 9 mice (het-cKO, GBR12909), and *n* = 14 from *N* = 8 mice (het-cKO, SCH23390). \**p* < 0.05; \*\**p* < 0.01; \*\*\**p* < 0.001. Transient comparisons were Holm-adjusted; lick-frequency comparisons required no multiplicity adjustment.

### Signal analysis

Fluorescence traces were processed separately by region using dual-wavelength reference correction and biexponential detrending [24–27]. Slow 405- and 470-nm baseline components were estimated with a 0.1-Hz low-pass filter and biexponential fits; the 405-nm reference was scaled by the fitted baseline ratio before subtraction. Signals were z-scored using pooled inter-trial interval (ITI) samples, retaining the corresponding baseline-recording SD for paired pharmacological comparisons [12, 28]. Outcome- and CS-evoked amplitudes were the first local maxima within 0–3 s after first lick and 0–1.5 s after CS onset, respectively; duration was quantified as peak width at half maximum (half-width) [28].

Spontaneous ITI transients were detected from detrended and smoothed signals using *z* > 1.645, minimum width 0.5 s, and minimum separation 2 s [28, 29]. Frequency was expressed as events/min and duration as half-width. Spectral organization was calculated from the full 40-Hz z-scored trace by FFT, integrated over six predefined bands from 0.01 to 10 Hz, and log10-transformed [13, 14]. Trial-history analysis used nonoverlapping Air-puff→Reward and Omission→Reward pairs and session-level mean CS amplitudes. Full preprocessing and analysis procedures are provided in the Supplementary Materials and Methods.

### DAT analysis

DAT expression was assessed in a separate cohort by western blotting in the caudate– putamen (CPu), NAc, TS, and cerebellum (Cb) and normalized to β-actin. The specificity of the anti-DAT monoclonal antibody used in this study has been evaluated previously [30].

### Statistics

Analyses were performed in MATLAB R2026a (MathWorks, Natick, MA, USA). All tests were two-tailed. Omnibus fixed-effect *p* values were unadjusted; *p* < 0.05 was considered significant. Prespecified contrasts were Holm-adjusted within defined comparison families; *p*_Holm_ < 0.05 was considered significant and 0.05 ≤ *p*_Holm_ < 0.10 a trend. LMEs were fit by restricted maximum likelihood with Satterthwaite degrees of freedom. Principal session-level LME analyses for Figs. 3d–j and 4 were independently repeated in R 4.6.1 (R Foundation for Statistical Computing, Vienna, Austria), yielding essentially identical estimates and *p* values (Supplementary Materials and Methods).

**Figure 3.**
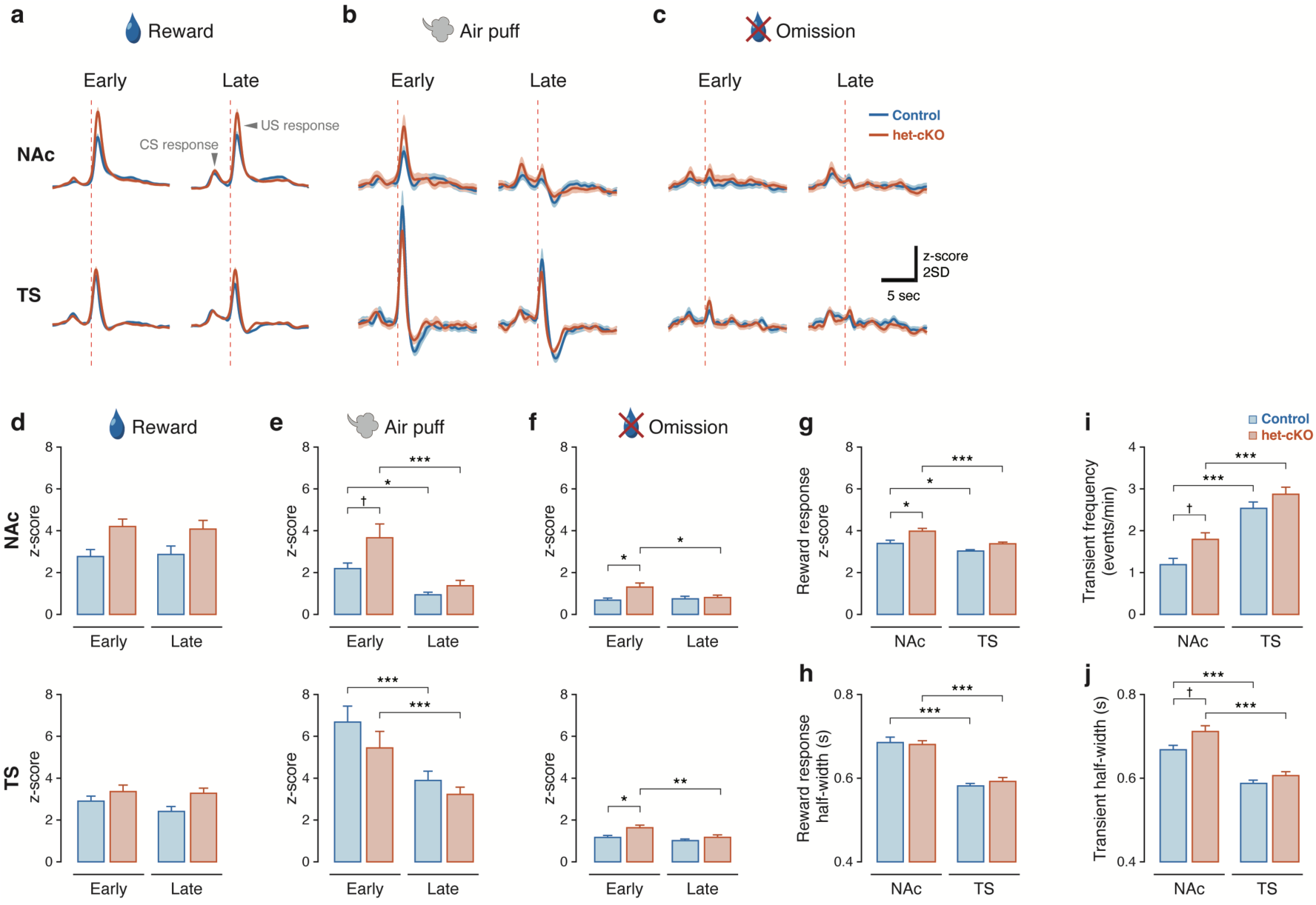
Outcome-evoked and spontaneous striatal dopamine dynamics. (a–c) Dopamine responses aligned to the first lick during the response period in Reward (a), Air-puff (b), and Omission trials (c), separated by Region, Genotype, and Phase. Traces show means and 95% confidence intervals across sessions. (d–f) US-response amplitudes for Reward (d), Air-puff (e), and Omission (f), defined as the first local maximum within 0–3 s after first-lick onset. NAc and TS are shown above and below, respectively. Each Region and Outcome was analyzed using a session-level Phase × Genotype linear mixed-effects model with Holm-adjusted comparisons (*n* = 20 sessions from *N* = 10 mice in each Genotype × Phase combination). (g, h) Reward-response amplitude (g) and half-width (h), pooled across drug-free Test sessions from day 8 onward. (i, j) Frequency (i) and half-width (j) of spontaneous ITI transients. Data in (g–j) were compared by Region and Genotype using session-level mixed-effects models with Holm-adjusted contrasts. Paired NAc–TS recordings: control, up to *n* = 70 sessions; het-cKO, up to *n* = 68 sessions; *N* = 10 mice per genotype. Bars show session means with upper SEM error bars. \**p* < 0.05; \*\**p* < 0.01; \*\*\**p* < 0.001; †0.05 ≤ *p* < 0.1 (Holm-adjusted).

**Figure 4.**
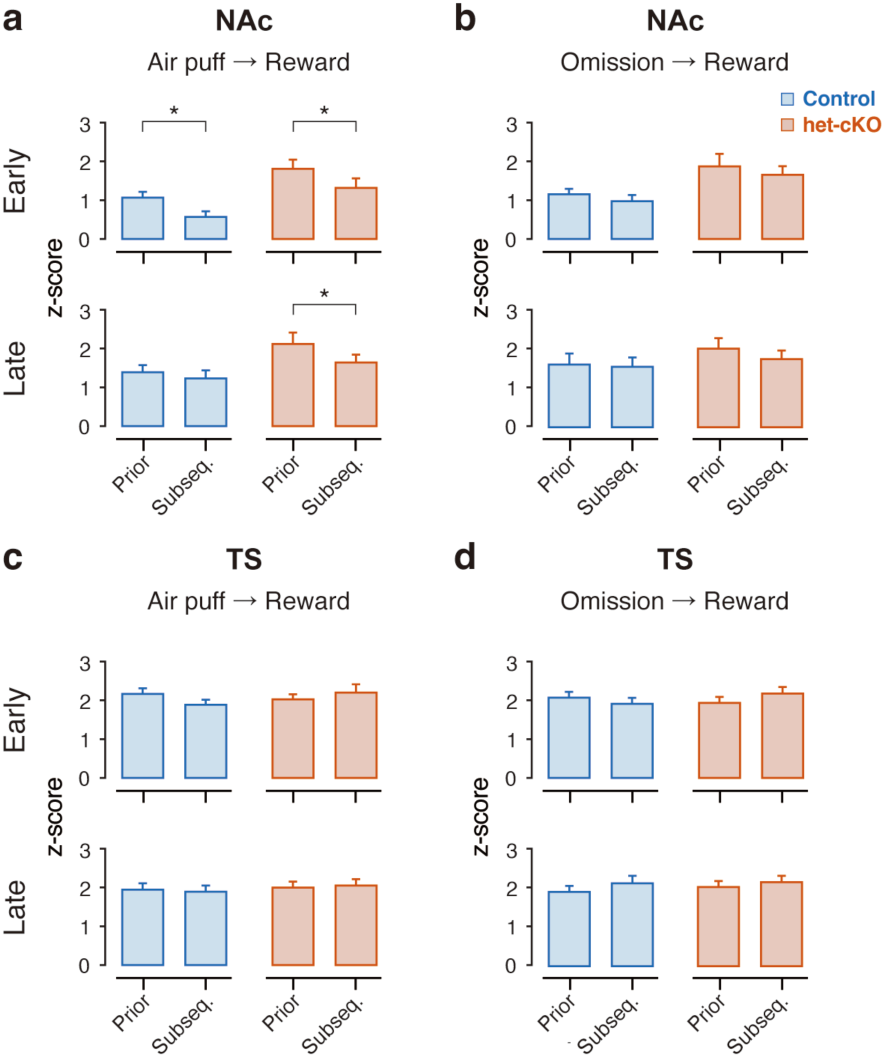
Trial-history-dependent modulation of CS-evoked dopamine responses. Consecutive, nonoverlapping trial pairs were classified as Air-puff→Reward or Omission→Reward. CS-response amplitude was defined as the first local maximum within 0–1.5 s after CS onset. Prior and Subseq. denote the preceding Air-puff or Omission trial and the subsequent Reward trial, respectively. (a) NAc, Air-puff→Reward. (b) NAc, Omission→Reward. (c) TS, Air-puff→Reward. (d) TS, Omission→Reward. Responses were averaged across all corresponding nonoverlapping trial pairs within each session and analyzed separately for each Region, Outcome, and Phase using Sequence × Genotype linear mixed-effects models with Holm-adjusted Prior– Subsequent comparisons. Bars show session means with upper SEM error bars (*n* = 20 sessions from *N* = 10 mice in each Genotype × Phase combination). \**p* < 0.05 (Holm-adjusted).

Males and females were pooled in the primary analyses; exploratory sex-sensitivity analyses are summarized in Supplementary Table S5.

For Fig. 1f,g, two sessions per phase were averaged within mouse and analyzed by Genotype × Phase mixed-design repeated-measures ANOVA. Mauchly’s test indicated violations of sphericity for both Fig. 1f (*W* = 0.0778, *ε*_GG_ = 0.451) and Fig. 1g (*W* = 0.356, *ε*_GG_ = 0.608); Greenhouse–Geisser corrections were therefore applied to Phase-containing effects. For Fig. 2b, session spectra were averaged within mouse, and bandwise genotype differences were tested by Welch’s *t* tests with Holm correction within region. DAT expression in Fig. 5c was compared between genotypes within each region by Welch’s *t* tests with Holm correction across four regions.

**Figure 5.**
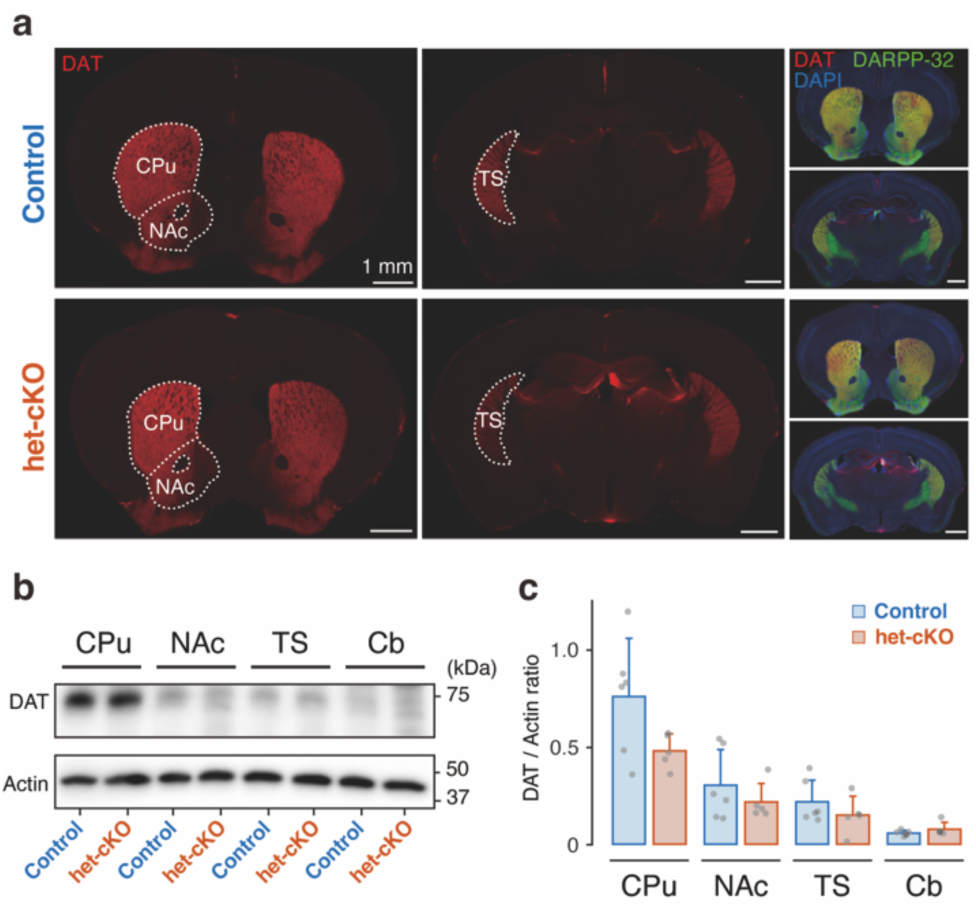
Regional dopamine transporter expression in control and het-cKO mice. (a) Representative coronal sections showing dopamine transporter (DAT) immunoreactivity. Dotted outlines indicate the CPu, NAc, and TS. Merged images show DAT (red), DARPP-32 (green), and DAPI (blue). Scale bars, 1 mm. (b) Representative western blots of DAT and β-actin in the CPu, NAc, TS, and cerebellum (Cb). (c) DAT expression normalized to β-actin. Bars show means with upper SD error bars across mice; gray points represent individual mice (control, *N* = 6; het-cKO, *N* = 5). Genotypes were compared within each region using Welch’s *t* tests with Holm correction across the four regions.

For the phase-independent analyses in Fig. 3g–j, session-level data from days 8, 9, 11, 12, 15, 17, and 19 were pooled; sessions from day 12 onward were recorded before drug administration, and post-drug sessions were excluded. The day 12 recording served both as the second Late-phase session and as the Before-drug session and was included only once. Because the Early- and Late-phase sessions also contributed to the phase-specific analyses in Fig. 3d–f, the datasets underlying Figs. 3d–f and 3g–j partially overlapped and should not be regarded as independent replications.

Session-level LMEs were Outcome ∼ Treatment + (1|MouseID) + (1|PairID) for complete Before–After pharmacological session pairs in Fig. 2d–f, USResponse ∼ Phase × Genotype + (1|MouseID) for Fig. 3d–f, MetricValue ∼ Region × Genotype + (1|MouseID) + (1|SessionID) for Fig. 3g–j, and CSResponse ∼ Sequence × Genotype + (1|MouseID) + (1|SessionID) for Fig. 4. Prespecified comparisons were Holm-adjusted by family. Effect estimates and unadjusted 95% confidence intervals (CIs) for key prespecified contrasts are reported in Supplementary Table S6; multiplicity-adjusted inference was based on Holm-adjusted *p* values. The mouse was the statistical unit for behavioral analyses, mouse-level spectral analyses, and DAT expression. Complete Before–After session pairs and individual recording sessions were the observational units for the remaining photometry analyses, with repeated observations accounted for by MouseID and, where applicable, PairID or SessionID random effects.

## Results

### *Atp2a2* het-cKO mice acquired the conditioning task comparably to controls

Histology confirmed dLight1.2 expression and fiber placement in the NAc and TS (Fig. 1a). Representative ATP2A2 western blots are shown in Fig. 1b; brain-specific ATP2A2 reduction in this line was established previously [10]. Both groups progressively acquired task-related licking (Fig. 1e).

Response-period Δlick frequency showed a significant Phase effect (*F*(1.35, 24.36) = 27.29, *p*_GG_ = 5.04 × 10^−6^), but no Genotype effect or Genotype × Phase interaction (*p* = 0.286 and *p*_GG_ = 0.402; Fig. 1f). Holm-adjusted comparisons showed increases across training in both genotypes, whereas Training-to-Test Early was nonsignificant in either group and no between-genotype comparison was significant.

Anticipatory Δlick frequency likewise showed a Phase effect (*F*(1.22, 21.90) = 23.09, *p*_GG_ = 3.82 × 10^−5^), but no Genotype effect or interaction (*p* = 0.850 and *p*_GG_ = 0.932; Fig. 1g). Thus, conditioning developed comparably between genotypes.

### The fiber-photometry system captured region-specific and pharmacologically sensitive dopamine dynamics in the striatum

Simultaneous NAc and TS recordings showed outcome-related dopamine fluctuations during Reward, Air-puff, and Omission trials (Fig. 2a), and Fig. 2c summarizes the pharmacological schedule.

Mouse-level spectral power did not differ between genotypes in either region (NAc, all *p*_Holm_ ≥ 0.200; TS, all *p*_Holm_ ≥ 0.838; Fig. 2b). Complementary analyses showed region-dependent spectral organization, no genotype difference in GBR-induced spectral changes, and no residual spectral difference in recovery recordings (Fig. S1).

GBR12909 increased NAc ITI transient frequency and half-width in both genotypes (all *p*_Holm_ ≤ 0.0155), whereas SCH23390 decreased frequency in both groups (both *p*_Holm_ ≤ 0.00139) and shortened half-width only in het-cKO mice (*p*_Holm_ = 0.0446; Fig. 2d,e). Licking increased after GBR12909 and decreased after SCH23390 in both genotypes (all *p* ≤ 0.00204; Fig. 2f); corresponding TS results are shown in Fig. S2. GBR12909 provided the primary pharmacological validation of dopamine sensitivity. Because SCH23390 can directly antagonize the D1-receptor-derived dLight1.2 sensor, its photometric effects were not interpreted as independent evidence of dopamine specificity; the concomitant licking reduction confirmed a systemic pharmacological effect.

### *Atp2a2* haploinsufficiency enhanced rapid dopamine signaling in an outcome- and region-dependent manner

Trial-averaged traces showed distinct Reward-, Air-puff-, and Omission-related responses in the NAc and TS during Early and Late phases (Fig. 3a–c).

Outcome-evoked dopamine responses showed strong learning-related attenuation to air puff and genotype-associated enhancement most clearly for omission. Reward responses showed no significant planned comparison; the NAc Genotype effect did not reach statistical significance (*F*(1, 18) = 4.20, *p* = 0.0553), without a Phase × Genotype interaction (*p* = 0.648; Fig. 3d). Air-puff responses decreased markedly from Early to Late in both regions (NAc, *F*(1, 58) = 35.03, *p* = 1.86 × 10^−7^; TS, *F*(1, 58) = 43.50, *p* = 1.39 × 10^−8^). The decrease was significant in both genotypes in both regions (NAc: control, *p*_Holm_ = 0.0133; het-cKO, *p*_Holm_ = 5.03 × 10^−6^; TS: control, *p*_Holm_ = 6.49 × 10^−6^; het-cKO, *p*_Holm_ = 5.67 × 10^−4^). In the NAc, the planned Early genotype contrast showed a trend toward a higher response in het-cKO mice (*p*_Holm_ = 0.0650), whereas the Phase × Genotype interaction did not reach statistical significance (*F*(1, 58) = 2.99, *p* = 0.0890; Fig. 3e).

Genotype-related enhancement was strongest for responses to reward omission during the Early phase. In the NAc, omission responses showed a significant Genotype effect (*F*(1, 18) = 4.94, *p* = 0.0393) and Phase × Genotype interaction (*F*(1, 58) = 4.59, *p* = 0.0365). Responses were greater in het-cKO mice during Early (*p*_Holm_ = 0.0140) and decreased from Early to Late only in het-cKO mice (*p*_Holm_ = 0.0277). In the TS, omission responses showed significant Phase (*F*(1, 58) = 9.52, *p* = 0.00312) and Genotype effects (*F*(1, 18) = 6.67, *p* = 0.0188), with a larger Early response in het-cKO mice (*p*_Holm_ = 0.0143) and an Early-to-Late decrease in that genotype (*p*_Holm_ = 0.00724). The Phase × Genotype interaction was not significant (*p* = 0.129; Fig. 3f). Direct regional comparisons are shown in Fig. S3.

Across regions, reward responses showed pronounced spatial organization together with a modest genotype-associated increase in amplitude. Reward-response amplitude showed significant Region (*F*(1, 252.92) = 25.09, *p* = 1.03 × 10^−6^) and Genotype effects (*F*(1, 17.82) = 4.46, *p* = 0.0491). Amplitude was higher in het-cKO mice in the NAc (*p*_Holm_ = 0.0469; estimated het-cKO − control difference = 0.687 z-score, 95% CI 0.102–1.273) and higher in the NAc than TS in controls (*p*_Holm_ = 0.0273) and het-cKO mice (*p*_Holm_ = 5.27 × 10^−5^; Fig. 3g). Reward-response half-width showed a strong Region effect (*F*(1, 252.64) = 136.04, *p* = 1.96 × 10^−25^) and was longer in the NAc in both genotypes (both *p*_Holm_ ≤ 3.36 × 10^−12^), whereas neither the Genotype effect nor the Region × Genotype interaction was significant (Fig. 3h).

Spontaneous ITI transients showed a complementary regional organization, with higher event frequency in the TS and longer duration in the NAc, while genotype-related increases in NAc transient measures remained at trend level. Transient frequency showed a significant Region effect (*F*(1, 253.74) = 79.27, *p* = 1.06 × 10^−16^) and was higher in the TS in both genotypes (both *p*_Holm_ ≤ 2.11 × 10^−7^). The Genotype effect did not reach statistical significance (*F*(1, 17.63) = 3.45, *p* = 0.0799), whereas the planned NAc genotype contrast showed a trend toward higher frequency in het-cKO mice (*p*_Holm_ = 0.0950; Fig. 3i). Transient half-width likewise showed a significant Region effect (*F*(1, 245.51) = 80.50, *p* = 7.70 × 10^−17^) and was longer in the NAc in both genotypes (both *p*_Holm_ ≤ 4.00 × 10^−7^). The Genotype effect did not reach statistical significance (*F*(1, 16.32) = 4.37, *p* = 0.0527), whereas the planned NAc genotype contrast showed a trend toward a greater transient half-width in het-cKO mice (*p*_Holm_ = 0.0520; estimated het-cKO − control difference = 0.0421 s, 95% CI 0.0053–0.0788; Fig. 3j). No Region × Genotype interaction was significant for either transient metric.

### Aversive trial history modulated cue-evoked dopamine signaling in the NAc

Recent aversive history modulated NAc cue-evoked dopamine responses, with a Late-phase Prior–Subsequent difference detectable only in het-cKO mice, although the genotype dependence of this persistence was not statistically established. In the NAc Early Air-puff→Reward sequence, significant Sequence (*F*(1, 38) = 13.25, *p* = 8.07 × 10^−4^) and Genotype effects (*F*(1, 18) = 5.43, *p* = 0.0316) were detected, with significant Prior–Subsequent differences in both genotypes (control: estimated Prior − Subsequent difference = 0.496 z-score, 95% CI 0.109–0.883; het-cKO: 0.489 z-score, 95% CI 0.102– 0.876; both *p*_Holm_ = 0.0269; Fig. 4a). In Late, the Sequence effect remained significant (*F*(1, 38) = 7.70, *p* = 0.00851), and the planned Prior–Subsequent difference was significant only in het-cKO mice (*p*_Holm_ = 0.0108; estimated Prior − Subsequent difference = 0.476 z-score, 95% CI 0.149–0.802), whereas the corresponding control contrast was not significant (estimated difference = 0.157 z-score, 95% CI −0.169–0.483; *p*_Holm_ = 0.336). The Sequence × Genotype interaction was nonsignificant (*F*(1, 38) = 1.95, *p* = 0.170), precluding a direct genotype-difference claim.

NAc Omission→Reward sequences and all TS planned Prior–Subsequent comparisons were nonsignificant (Fig. 4b–d), and no Sequence × Genotype interaction reached significance. Thus, detectable Late-phase aversive-history modulation was confined to the NAc het-cKO group without statistical evidence for a genotype interaction.

### Regional DAT expression showed no statistically detectable genotype differences

DAT immunoreactivity was evident in the CPu, NAc, and TS in both genotypes (Fig. 5a), and representative western blots are shown in Fig. 5b. No genotype difference was detected in any region (all *p*_Holm_ = 1; Fig. 5c), and all mean-difference 95% CIs included zero. Effect sizes and CIs are reported in Supplementary Table S6.

## Discussion

Brain-specific Atp2a2 haploinsufficiency enhanced NAc reward-evoked dopamine responses in the pooled analysis, increased responses to unexpected reward omission in both regions during the Early phase, and was associated with trends toward increased spontaneous NAc transient frequency and half-width. Aversive trial history also continued to modulate subsequent NAc dopamine responses in het-cKO mice during Late, although a genotype difference in persistence was not established. These changes occurred despite comparable conditioning and no statistically detectable genotype difference in regional DAT expression.

Regional signal organization and pharmacological responses supported the validity of our custom-built fiber-photometry system. Genotypes did not differ in spectral power (Fig. 2b), whereas low-frequency power was greater in the NAc and high-frequency power in the TS (Fig. S1a,b), consistent with regional differences in dopamine timescales [13]. GBR12909 provided the principal pharmacological validation: DAT inhibition increased transient frequency and half-width, increased licking (Figs. 2d–f and S2), and shifted spectral power toward lower frequencies (Fig. S1c,d), consistent with previous dLight studies [12, 13, 31]. The larger TS spectral shift may reflect regional differences in dopamine uptake and clearance [14, 32, 33]. SCH23390 reduced photometric signals and licking, but because dLight1.2 is D1-receptor-derived and can be directly antagonized by SCH23390 [12, 31, 34], its photometric effects cannot independently establish reduced extracellular dopamine. Together, the GBR12909 response and regional signal characteristics support the ability of our system to monitor extracellular dopamine dynamics.

Training increased licking (Fig. 1f,g), reduced air-puff US responses in both regions (Fig. 3e), and increased the NAc CS response (Fig. S4a), consistent with transfer of dopamine signaling toward predictive cues [11, 35]. Related attenuation of aversive outcome-evoked TS dopamine has been reported during associative fear learning [18] and repeated air-puff exposure [36]. Regionally, reward responses were larger and longer in the NAc (Fig. 3g,h), whereas spontaneous transients were longer in the NAc but more frequent in the TS (Fig. 3i,j). Direct regional comparisons also showed larger air-puff responses in the TS across phases and genotypes (Fig. S3b). These patterns accord with established striatal gradients in dopamine timescales and preferential recruitment of posterior striatal dopamine by novel or threatening events [13–18]. The larger TS omission response in the direct regional analysis further suggests that this specialization may extend to unexpected reward omission (Fig. S3c).

The het-cKO mice showed a modest, context-dependent enhancement of short-timescale dopamine signals. In the pooled analysis, reward-response amplitude was higher in het-cKO mice in the NAc (Fig. 3g), while NAc spontaneous transient frequency and half-width showed trends toward increases (Fig. 3i,j). In the phase-specific analysis, the Early NAc air-puff response also showed a trend toward enhancement (Fig. 3e). Omission responses were greater in het-cKO mice during Early in both regions, with a significant Phase × Genotype interaction in the NAc but not the TS (Fig. 3f). Thus, *Atp2a2* deficiency appears to amplify rapid dopamine signaling under specific regional and experiential conditions rather than producing a uniform elevation.

Recent aversive history continued to suppress subsequent reward-predictive CS responses in the NAc during Late only in het-cKO mice (Fig. 4a), although the Sequence × Genotype interaction was nonsignificant. No comparable Omission→Reward or TS history effect was detected (Fig. 4b–d), arguing against generalized cue suppression. This pattern may reflect latent-state inference or persistent aversive salience [37, 38] and, together with impaired fear memory in het-cKO mice [10], suggests altered temporal processing of aversive information. Similar abnormalities in ventral-striatal responses and salience attribution occur in schizophrenia [39–41]. Determining whether these parallels reflect shared alterations in latent-state inference or salience processing will require tasks that dissociate these processes more directly.

To investigate the underlying mechanism, we next asked whether impaired dopamine clearance explained these phenotypes. Prior microdialysis showed elevated extracellular NAc dopamine in het-cKO mice [10]. In our system, GBR12909 prolonged transient half-width in both genotypes and regions (Figs. 2e and S2), confirming that reduced DAT-mediated clearance lengthens dLight signals. Under drug-free conditions, reward-response half-width did not differ by genotype (Fig. 3h), and only NAc transient half-width showed a trend toward prolongation (Fig. 3j). No statistically detectable genotype difference in DAT protein expression was observed in any of the four regions examined (Fig. 5c), although the confidence intervals were sufficiently wide that equivalence between genotypes cannot be established. The ER-resident redox protein SELENOT interacts with ATP2A2 to regulate Ca^2+^ flux and NURR1-dependent DAT expression, and *Selenot* deletion causes hyperdopaminergia with reduced DAT expression [42]. The absence of a statistically detectable reduction in regional DAT expression in our model therefore differs from the DAT reduction reported after SELENOT loss. Because major DAT impairment typically prolongs extracellular dopamine signals [31, 43], the present findings do not support a major reduction in DAT expression as the primary explanation for the dopamine phenotype. However, the relatively wide confidence intervals for DAT expression, together with the absence of direct uptake measurements, leave open the possibility of more subtle changes in DAT expression, trafficking, or uptake kinetics.

Altered dopamine-neuron activity or presynaptic release could contribute to the observed phenotype. ATP2A2-mediated Ca^2+^ transport into the ER limits residual cytosolic Ca^2+^, a key regulator of neurotransmitter release [44–46], and ATP2A2 knockdown in the NAc increases local dopamine and methamphetamine-conditioned place preference, whereas overexpression has opposite effects [47]. That abnormalities were more evident in evoked than spontaneous signals is consistent with an activity-dependent Ca^2+^-related mechanism. Thus, ATP2A2-dependent intracellular Ca^2+^ regulation represents one possible contributor to the altered dopamine signals and aversive-history effects observed here, a possibility that requires direct experimental testing.

This study has limitations. dLight photometry measures relative fluorescence and cannot separate release, axonal activity, diffusion, and clearance. Presynaptic Ca^2+^ and DAT uptake kinetics were not measured directly, and region was confounded with hemisphere because recordings were consistently obtained from the left NAc and right TS. Air-puff and omission trials were limited to five per session, restricting the number of nonoverlapping pairs available for history analysis. The modest cohort limited precision; several genotype-related findings remained at trend level and were attenuated when sessions were averaged within mouse, while sex-specific inference was underpowered. The broad age range may have added biological variability, and the history analysis cannot distinguish latent-state inference from persistent aversive salience or altered expectation. Larger cohorts, direct measurements of presynaptic Ca^2+^ dynamics, dopamine release, and uptake kinetics, and tasks manipulating state transitions are needed.

In summary, *Atp2a2* haploinsufficiency altered the magnitude, regional and temporal organization, and recent-outcome dependence of striatal dopamine signals without impairing conditioning or producing a statistically detectable reduction in regional DAT expression. These context-dependent alterations in striatal dopamine signaling may be relevant to the psychiatric vulnerability associated with *ATP2A2* loss-of-function, although the contributions of altered intracellular Ca^2+^ regulation and changes in dopamine release to these phenotypes remain to be established directly.

## Data Availability Statement

The custom MATLAB code used for data processing and statistical analysis will be made publicly available in a GitHub repository upon publication. The data that support the findings of this study are available from the corresponding authors upon reasonable request.

## Acknowledgments

We thank Dr. Tadafumi Kato and the RIKEN Center for Brain Science for sharing the *Atp2a2* het-cKO mice.

## Author Contributions

M.N.: Methodology, Investigation, Formal analysis, Data curation, Visualization, Writing – original draft.

T.Y.: Conceptualization, Methodology, Software, Investigation, Formal analysis, Data curation, Visualization, Supervision, Project administration, Funding acquisition, Writing – original draft, Writing – review & editing.

N.I.: Methodology, Investigation, Writing – review & editing. M.M.: Supervision, Writing – review & editing.

K.N.: Conceptualization, Resources, Methodology, Supervision, Project administration, Funding acquisition, Writing – original draft, Writing – review & editing.

T.H.: Conceptualization, Resources, Methodology, Supervision, Project administration, Funding acquisition, Writing – review & editing.

All authors approved the final version of the manuscript and agree to be accountable for all aspects of the work.

