## Supplementary information for "Striatal dopamine dynamics across temporal scales in brain-specific *Atp2a2* heterozygous knockout mice during reward conditioning"

Niido *et al.*

#### Supplementary Materials and Methods

*Reference citations in this Supplementary Information correspond to the reference list in the Main Manuscript.*

##### Animals, breeding, and genotyping

The *Atp2a2* conditional knockout line, generated as described previously (Nakajima *et al.*, 2021), was maintained on a C57BL/6N background.

Fiber-photometry experiments included 10 control mice (4 males and 6 females) and 10 het-cKO mice (5 males and 5 females), aged 16.6–50.4 weeks at the first Early-phase recording (Supplementary Table S2). Mice were housed in standard polyolefin cages (211 × 319 × 133 mm; CL-0104-2, CLEA Japan, Tokyo, Japan) under a reversed 12-h light/dark cycle (lights on, 10:00–22:00) at 24 °C and 50 ± 10% humidity. They were group-housed (2–5 mice per cage) before surgery and singly housed thereafter. During water restriction, body weight was monitored daily and maintained at ≥90% of the baseline value measured under *ad libitum* conditions. Birth, weaning, identification, and group allocation were pseudorandomized while balancing littermates.

All procedures were approved by the Animal Care and Use Committee of Teikyo University School of Medicine (No. 24-018) and the Teikyo University Genetically Modified Organism Experimental Safety Committee (No. 2404382A1) and were conducted in accordance with the approved guidelines.

##### Stereotaxic surgery

Mice were anesthetized intraperitoneally with medetomidine hydrochloride (0.75 mg/kg), midazolam (4 mg/kg), and butorphanol tartrate (5 mg/kg) dissolved in saline and positioned in a stereotaxic apparatus. pAAV-hSyn-dLight1.2 (AAV5; 1.0 × $10^{13}$ genome copies/mL; Addgene #111068-AAV5) was injected unilaterally into the left nucleus accumbens (NAc; anteroposterior +1.2 mm, mediolateral +1.0 mm, dorsoventral −4.5 mm from bregma) and right tail of the striatum (TS; anteroposterior −1.0 mm, mediolateral −3.25 mm, dorsoventral −2.5 mm from bregma). TS coordinates were selected with reference to Willmore *et al.* (2022). The dLight1 sensor family was originally validated by Patriarchi *et al.* (2018).

At each site, 1.0 µL of vector was delivered at 100 nL/min and 2.0 psi through a glass micropipette (30–50-µm tip) connected to a FemtoJet 5247 microinjector (Eppendorf, Hamburg, Germany). The pipette remained in place for 15 min and was then withdrawn gradually over approximately 5 min to minimize backflow.

An optical-fiber cannula (400-µm core, 0.39 numerical aperture; CFML14L05, Thorlabs, or R-FOC-BL400C-39NA, RWD) was implanted immediately above each injection site and secured with Super-Bond C&B dental resin (Sun Medical). A custom stainless-steel head plate was attached during the same procedure. Behavioral training and recording began after at least 2 weeks to permit AAV-mediated sensor expression.

##### Behavioral apparatus, schedule, and analysis

Experiments used a head-fixed classical conditioning task adapted from published appetitive and aversive conditioning procedures (Deseyve *et al.*, 2024; Walle *et al.*, 2024). All experiments were conducted in a darkened, sound-attenuating chamber. The conditioned stimulus (CS) consisted of a 10-kHz pure tone amplitude-modulated at 10 Hz together with a blue LED flashing at 10 Hz (irradiance, 1.0 mW/m$^2$). Reward was approximately 3–5 µL of 0.1% saccharin delivered through a spout by a tube pump (RP-HX01S-1A-DC3VS; Aquatech, Osaka, Japan), whereas the aversive outcome was a 0.5-s air puff directed toward the nose using a vacuum pump (ROB-10398; SparkFun Electronics, Niwot, CO, USA). Licks were detected by interruption of an infrared beam (E3Z-G61-M3J; Omron, Kyoto, Japan), and the first interruption after response-period onset was defined as first-lick onset.

For frontal video recording, the mouse's face was illuminated with an infrared LED ring (FRS5CC; OptoSupply, Hong Kong). Video was acquired with a DMK33UX273 camera (The Imaging Source, Bremen, Germany) at 40 fps with an exposure time of 1/154 s, 320 × 200-pixel resolution after 2 × 2 binning, 4.8-dB gain, and 8-bit depth. Behavioral events, photometry frames, and frontal video were synchronized using shared TTL timing signals and custom MATLAB scripts.

The schedule comprised Pre-training on days 1–3, Training on days 4–5, rest on days 6–7, and Test on days 8–12. For phase-based analyses, days 8 and 9 were defined as the Early phase and days 11 and 12 as the Late phase. Day 10 was an intervening Test session and was not included in the phase-based analyses. Pre-training sessions contained 200 trials; the CS/response period and intertrial interval (ITI) were each 3 s, and the first lick during the CS triggered reward. Training sessions contained 150 trials; the 6-s CS comprised a 3-s delay period followed by a 3-s response period, the first response-period lick triggered reward, and the ITI was 6 s. Test sessions contained 50 trials; the 10-s CS comprised a 3-s delay period followed by a 7-s response period, and the ITI varied pseudorandomly from 20 to 35 s. Reward, air-puff, and omission trials occurred in a pseudorandom order at session-level frequencies of 80%, 10%, and 10%, respectively. During Test, the first response-period lick triggered the assigned outcome. Each Test session was followed by one 150-trial Training session except on day 12. Complete phase parameters are listed in Supplementary Table S3.

Data analysis was performed in MATLAB R2026a (MathWorks, Natick, MA, USA). Lick peri-stimulus time histograms were aligned to CS onset. Licks were counted in nonoverlapping 50-ms bins, averaged across trials within session, and converted to licks/s without temporal smoothing. The baseline-subtracted lick frequency (Δlick frequency) was the mean in the analysis window minus the mean from −2 to 0 s relative to CS onset. Response-period licking was quantified during the first 2 s after response-period onset. Anticipatory licking was quantified during 1–3 s after CS onset, a window common to Training and Test.

##### Drug preparation and administration

GBR12909 dihydrochloride (Tocris Bioscience, #0421/10) and SCH23390 hydrochloride (MedChemExpress, HY-19545A) were dissolved in physiological saline immediately before use. GBR12909 was used to inhibit the dopamine transporter (DAT; Holloway *et al.*, 2023), whereas SCH23390 was used to block D1 receptors (Patriarchi *et al.*, 2018). GBR12909 (20 mg/kg, intraperitoneal) and SCH23390 (0.25 mg/kg, intraperitoneal) were administered after a 50-trial Before-drug Test session. A second 50-trial Test session was conducted 60 min after administration (After-drug). GBR12909 was administered on days 12 and 17 and SCH23390 on days 15 and 19. The day 12 Before-drug recording was identical to the second Late-phase Test session. For the recovery analyses shown in Figs. S1e,f, the pre-drug recordings on days 15 and 19 were used as recovery recordings after the preceding GBR12909 administration, whereas the pre-drug recording on day 17 was used as the recovery recording after the preceding SCH23390 administration. Separate drug-free Test sessions were also performed on days 16 and 18, but these were not used for the recovery analyses shown in Figs. S1e,f; each was followed by a 150-trial Training session. Experimental-day assignments are summarized in Supplementary Table S4.

#### Dual-site fiber photometry system

A camera-based dual-site fiber-photometry system was constructed with reference to the multi-fiber architecture described by Dai *et al.* (2024). dLight1.2 was excited alternately with 405-nm (M405FP1, Thorlabs) and 470-nm (M470F4, Thorlabs) LEDs at 40 Hz per wavelength. The LEDs were driven by LEDD1B LED drivers (Thorlabs). Excitation light was filtered with FBH405-10 and MF469-35 bandpass filters (Thorlabs), combined by an MD416 dichroic mirror, reflected by a DMLP490 long-pass dichroic, and delivered through a PLN 10× objective (Olympus) and a bifurcated low-autofluorescence fiber bundle (BBP(2)_400/440/900-0.37_2m_FCM-2xMF1.25_LAF; Doric Lenses).

Excitation light power was measured with a PM160 optical power meter (Thorlabs) at the distal tip of an optical fiber identical to the implanted fibers and connected through the same patch-cord and mating-sleeve configuration used for recordings. Under continuous illumination, the 470-nm excitation power was 460 μW for the NAc channel and 309 μW for the TS channel, whereas the 405-nm excitation power was 126 and 81.0 μW, respectively. During the interleaved acquisition mode used for recordings, the corresponding time-averaged powers were 218 and 146.5 μW at 470 nm and 59.8 and 39.4 μW at 405 nm for the NAc and TS channels, respectively. The lower powers measured during interleaved illumination reflected the duty cycle of the alternating excitation sequence.

Emitted fluorescence returned through the same optical path, passed through the DMLP490 dichroic, was filtered at 525 nm (MF525-39, Thorlabs), and was imaged with a DMK33UX273 CMOS camera using IC Express (The Imaging Source). Output faces of the NAc and TS patch cords were imaged simultaneously on the same sensor. Photometry images were acquired at 80 fps with an exposure time of 1/92 s, a resolution of 400 × 200 pixels after 2 × 2 binning, 8-bit depth, and a gain of 20.2–27.8 dB. LED illumination, photometry-camera frames, face-camera frames, behavioral events, and video were synchronized by TTL signals generated by a custom Arduino-based controller (Yoshida, 2026).

#### Image preprocessing and temporal alignment

Signal processing was performed in MATLAB R2026a using custom scripts. Frames acquired during alternating 405- and 470-nm illumination were initially separated according to the acquisition sequence. Frame timestamps were then inspected for temporal discontinuities or dropped frames. Such inconsistencies were rare; when detected, a uniformly sampled 40-Hz time axis was reconstructed, and each time point was assigned the recorded frame with the nearest timestamp. Thus, occasional missing frames were handled by temporal reassignment of recorded frames rather than by image interpolation. Recording onset differences among simultaneously acquired camera streams were also corrected on the basis of their timestamps, and temporally aligned image sequences were generated for subsequent analysis.

Task-event timing was independently obtained from the synchronized face-camera recording. Separate region of interests (ROIs) were manually defined over the lick-related image region and the task LED, and the mean pixel intensity within each ROI was extracted frame by frame. CS onset and offset were initially detected from changes in the LED-intensity signal. When reliable detection could not be achieved from intensity changes alone, the LED trace was analyzed in the time-frequency domain, and power around the 10-Hz flicker frequency was used to recover stimulus timing.

Detected CS events were matched to the behavioral task log. Behavioral timestamps were transformed onto the video/photometry time axis using trial-by-trial temporal scaling based on consecutive CS onsets, thereby correcting small timing differences and accumulated clock drift between acquisition systems. The response period was defined as the interval from 3 s after CS onset to CS offset, and the first lick occurring within this period was designated the first lick. Inter-trial intervals (ITIs) were defined as the periods from CS offset to the onset of the subsequent trial.

##### ROI definition and extraction of fluorescence traces

A circular ROI was manually placed over each fiber output face on a reference video frame. The ROIs were converted into binary masks and retained for subsequent analysis. The mean pixel intensity within each ROI was then calculated for every frame, yielding one NAc and one TS fluorescence trace for each excitation series.

The two excitation series were subsequently assigned to the 405- and 470-nm channels on the basis of their frequency-domain signal characteristics. The series exhibiting the greater overall spectral magnitude was assigned to the 470-nm excitation channel, with the other series assigned to the 405-nm channel.

##### Fluorescence preprocessing

Each 40-Hz trace was processed separately for each recording region. Slow baseline components were isolated from the 470- and 405-nm traces using a sixth-order Butterworth low-pass filter with a cutoff frequency of 0.1 Hz, and a biexponential function was fitted to each low-pass-filtered trace to model slow drift and photobleaching:

$$B(t) = a\, e^{bt} + c\, e^{dt}$$

To account for time-dependent differences in baseline fluorescence between channels, the 405-nm trace was scaled pointwise according to the ratio of the fitted 470- and 405-nm baselines,

$$F_{405,\,\mathrm{scaled}}(t) = F_{405}(t) \times \frac{B_{470}(t)}{B_{405}(t)}$$

and subtracted from the 470-nm trace:

$$F_{\mathrm{diff}}(t) = F_{470}(t) - F_{405,\,\mathrm{scaled}}(t)\,.$$

The resulting difference signal was corrected for photobleaching by normalizing the fitted 470-nm baseline to its mean over the first 10 s,

$$L(t) = \frac{B_{470}(t)}{\langle B_{470}(0 - 10\ \mathrm{s})\rangle}$$

and calculating

$$F_{\mathrm{corr}}(t) = \frac{F_{\mathrm{diff}}(t)}{L(t)}\,.$$

This study-specific correction procedure combined established principles of dual-wavelength reference correction and exponential detrending used in fiber photometry. The 405-nm signal was used as an activity-independent reference for shared signal fluctuations (Lerner *et al.*, 2015; Kim *et al.*, 2016), while slow drift and photobleaching were modeled using exponential or biexponential functions (Patel *et al.*, 2020; Murphy *et al.*, 2023). These principles were integrated here by scaling the 405-nm trace pointwise according to the ratio of the fitted 470- and 405-nm baselines before subtraction from the 470-nm signal.

For z-score normalization, all ITI samples in each analyzed recording were pooled to calculate the mean corrected fluorescence level for that session. The SD used for normalization was determined in advance from all ITI samples in the corresponding baseline recording and retained for subsequent paired analyses. Each recording was therefore centered using its own ITI mean ($\mu_{\mathrm{ITI}}$) but scaled using the ITI-derived SD ($\sigma_{\mathrm{ITI}}$) from the corresponding baseline recording:

$$z(t) = \frac{F_{\mathrm{corr}}(t) - \mu_{\mathrm{ITI}}}{\sigma_{\mathrm{ITI}}}\,.$$

For baseline recordings themselves, their own ITI-derived SD was used. In pharmacological experiments, After-drug recordings were normalized using the ITI SD obtained from the immediately preceding Before-drug recording from the same mouse and region, so that post-treatment changes were expressed on the predrug variability scale rather than being renormalized by post-treatment variance. The same baseline-referenced normalization principle was used for task-evoked and spontaneous-transient analyses. The baseline-referenced normalization described above was implemented specifically for the present study, whereas the use of z-scored and event-aligned dLight signals was informed by previous dLight photometry studies (Patriarchi *et al.*, 2018; Robinson *et al.*, 2019).

##### Task-evoked responses

Task-evoked dopamine responses were analyzed from the baseline-referenced z-scored fluorescence traces. Outcome-evoked traces were extracted only from trials in which a first lick was detected during the response period, aligned to first-lick onset, and analyzed separately for Reward, Air-puff, and Omission trials. CS-aligned traces were generated independently using CS onset as time zero. Event-aligned traces were extracted over the predefined analysis window and smoothed using a 0.5-s moving-average filter.

Outcome-response amplitude was defined as the first local maximum within 0–3 s after first-lick onset. CS-response amplitude was defined as the first local maximum within 0–1.5 s after CS onset. Event-aligned responses were first averaged across the relevant trials within each session before subsequent session- or mouse-level analyses.

Response duration was quantified as peak width at half maximum (half-width). For each positive peak, the first half-height crossings on the rising and falling sides were identified, and the crossing times were estimated by linear interpolation between adjacent samples. Half-width was defined as the interval between the two interpolated crossings. Events for which both crossings could not be identified were treated as missing, and widths exceeding 3 s were excluded. Valid response widths were subsequently averaged within session. These peak- and half-width-based measures followed commonly used approaches for quantifying dLight responses (Robinson *et al.*, 2019).

##### Spontaneous transient analysis

Spontaneous dopamine transients were detected from the same baseline-referenced z-scored traces. Very slow components below 0.06 Hz were estimated using a sixth-order

Butterworth low-pass filter and subtracted from the z-scored trace. The residual signal was then smoothed using a 0.5-s moving-average filter.

Positive transients were identified from the processed z-scored trace using the MATLAB findpeaks function. Candidate peaks were required to exceed $z = 1.645$, an operational threshold approximately corresponding to the upper 5% of a standard normal distribution and comparable to the critical z-score of 1.645 previously applied to positive dLight dopamine responses (Kutlu *et al.*, 2022). Peaks were additionally required to have a minimum width of 0.5 s and to be separated from adjacent peaks by at least 2 s. Peaks occurring before the first CS onset or after the final CS offset were excluded. Among the remaining events, only peaks whose locations fell within ITIs, defined from CS offset to the subsequent CS onset, were included in the spontaneous-transient analysis. The use of threshold-based peak detection on z-scored dLight signals, including implementation with MATLAB findpeaks, was informed by previous dLight photometry analyses (Robinson *et al.*, 2019).

Transient amplitude was defined as the detected peak height. Transient frequency was calculated as the number of accepted ITI events divided by the total analyzed ITI duration and expressed as events/min. Transient duration was quantified as half-width using the same half-height crossing and linear-interpolation procedure described for task-evoked responses. Events lacking valid crossings were treated as missing, and widths exceeding 3 s were excluded. If no valid transient was detected in a recording, transient amplitude and half-width were treated as undefined, whereas transient frequency remained calculable from the total analyzed ITI duration.

##### Spectral analysis

Frequency-domain analysis used the full z-scored time series from each session and region; the signal was not restricted to ITIs. The analyzed spectral range was 0.01–20 Hz. For a trace $x$ of length $n$ sampled at 40 Hz, the discrete Fourier transform was computed with MATLAB fft, and power was calculated as $|\mathrm{FFT}(x)/n|^2$. The nonnegative-frequency portion from 0 to 20 Hz was retained, resampled to 10,000 equally spaced points, and smoothed with a 25-point moving average. Values that were nonfinite or ≤0 were replaced by machine epsilon. The 0-Hz component was excluded from all summaries.

Integrated power was calculated by summing spectral values within each of six half-open frequency bands—0.01–0.03, 0.03–0.1, 0.1–0.3, 0.3–1.0, 1.0–3.0, and 3.0–10 Hz—and multiplying the sum by the common frequency spacing. Band-power values were $\log_{10}$-transformed before statistical analysis. For mouse-level analyses, linear (untransformed) session spectra were first arithmetically averaged frequency-by-frequency within each MouseID; band integration and $\log_{10}$ transformation were then applied to the resulting mouse-level mean spectrum. Group spectral curves and 95% confidence intervals were calculated in $\log_{10}$ space and back-transformed for display.

Spectral analysis was used to characterize the temporal organization of dopamine signals across recording regions. This approach was motivated by previous frequency-domain analyses demonstrating region-dependent organization of dopamine dynamics (Jørgensen *et al.*, 2023), together with evidence that striatal dopamine signals vary across regions in their characteristic behavioral and reward-related timescales (Mohebi *et al.*, 2024). The specific frequency-band boundaries, integration procedure, and statistical transformations used here were defined for the present study.

#### Trial-history analysis

Within each Test session, consecutive trial pairs were classified as Air-puff→Reward or Omission→Reward. Pairs were selected without overlap, so that no trial contributed to more than one pair. For each sequence, CS-response amplitudes were calculated as the first local maximum within 0–1.5 s after CS onset and averaged separately across the preceding trial (Prior) and subsequent Reward trial (Subsequent). The two resulting session-level means formed a within-session paired observation. NAc and TS were processed separately, and Early and Late sessions were retained as separate phases.

#### Western blotting

Mice were euthanized by cervical dislocation, and brains were rapidly removed. The heart was also collected for ATP2A2 analysis to confirm the brain-specific reduction in ATP2A2 expression. Brains were cut into 1-mm coronal sections on ice, and the NAc, TS, caudate–putamen (CPu), and cerebellum (Cb) were dissected under a stereomicroscope. Each sample was homogenized in 50 mM Tris–HCl, 150 mM NaCl, 1% Triton X-100, 0.1% SDS, and Halt Protease and Phosphatase Inhibitor Cocktail, EDTA-free (100×; Thermo Fisher Scientific) diluted 1:100. Homogenates were incubated for 30 min at 4 °C, centrifuged at 15,000 rpm for 14 min at 4 °C, and the supernatants collected. Protein concentration was measured using the Pierce BCA Protein Assay Kit (Thermo Fisher Scientific, A65453) on a VersaMax microplate reader (Molecular Devices).

Samples were adjusted to 10 µg protein per lane and mixed with 4× Laemmli sample buffer (Bio-Rad, #1610747) containing 5% 2-mercaptoethanol. Aliquots for β-actin detection were heated at 95 °C for 5 min. ATP2A2 and DAT aliquots were not heated and were instead incubated at room temperature for 30 min. Proteins were separated on 4–20% Mini-PROTEAN TGX precast gels (Bio-Rad, #4561096) at a constant 200 V for approximately 60 min. Immobilon-P PVDF membranes (0.45-μm pore size; Merck Millipore, IPVH00010) were activated in methanol for 30 s and equilibrated in transfer buffer for 3 min before transfer at a constant 0.33 A for 3 h at 4 °C.

Membranes were washed in TBS-T (50 mM Tris–HCl, 150 mM NaCl, 0.1% Tween 20), blocked for 30 min at room temperature in TBS-T containing 2% skim milk, and incubated overnight at 4 °C with primary antibody in TBS-T containing 1% skim milk (1 mL per membrane). Primary antibodies were rabbit anti-ATP2A2/SERCA2 (Cell Signaling Technology, #4388; 1:400), mouse anti-DAT (mAb16; Thermo Fisher Scientific, MA5-24796; 1:400), and mouse anti-β-actin (clone AC-74; Sigma-Aldrich, A5316; 1:1,500). Membranes were washed three times for 5 min in TBS-T, incubated for 1 h at room temperature with HRP-conjugated donkey anti-rabbit IgG (Cytiva, NA934-1ML) or sheep anti-mouse IgG (Cytiva, NA931-1ML), each at 1:1,000, and washed three times for 10 min.

The anti-DAT monoclonal antibody used for both western blotting and immunohistochemistry in this study (mAb16; MA5-24796) has been evaluated previously for specificity (Russo *et al.*, 2023).

Chemiluminescence was developed using SuperSignal West Femto Maximum Sensitivity Substrate (Thermo Fisher Scientific, #34095) and acquired on an Amersham ImageQuant 680 system (Cytiva). Band intensities were quantified in Fiji/ImageJ 1.54p; each target band was normalized to β-actin from the same lane. Figure 1b shows representative ATP2A2 blots from one animal per genotype and was not used for quantitative group comparison. Reduced brain ATP2A2 expression in this het-cKO line was established quantitatively in the original characterization of the model [11]. DAT expression was quantified in a separate cohort from that used for fiber photometry, comprising control ($N$ = 6) and het-cKO mice ($N$ = 5), and is shown in Fig. 5b,c.

##### Immunohistochemistry

Mice were deeply anesthetized and perfused transcardially with 0.1 M phosphate-buffered saline (PBS), followed by 4% paraformaldehyde in PBS. Brains were removed, postfixed in 4% paraformaldehyde for 24 h at 4 °C, and cut into 50-μm coronal sections with a DTK-1000N microslicer (Dosaka EM). Free-floating sections were permeabilized for 30 min at room temperature in PBS containing 0.5% Triton X-100 and blocked for 1 h in PBS containing 10% normal goat serum and 0.1% Triton X-100. Primary antibodies diluted in blocking buffer were applied for 72 h at 4 °C with gentle agitation: rabbit anti-GFP (MBL, #598; 1:1,000), mouse anti-DAT (mAb16; Thermo Fisher Scientific, MA5-24796; 1:250), and rabbit anti-DARPP-32 (clone 19A3; Cell Signaling Technology, #2306; 1:400).

After three 10-min PBS washes, sections were incubated overnight at 4 °C with Alexa Fluor 488-conjugated goat anti-mouse IgG (Invitrogen, A11029; 1:200) and Alexa Fluor 594-conjugated goat anti-rabbit IgG (Invitrogen, A11037; 1:200), followed by PBS washes. Sections used to assess dLight1.2 expression and fiber placement were counterstained with NeuroTrace blue fluorescent Nissl stain (Invitrogen, N21479; 1:200); DAT/DARPP-32 sections were counterstained with DAPI (Dojindo, D523; 1:200). Sections were mounted in 45 mL glycerol plus 5 mL 1 M Tris–HCl and imaged as tiled fields with a BZ-X800 fluorescence microscope (Keyence, Osaka, Japan) and 4× objective. For presentation, DAT was pseudocolored red and DARPP-32 green.

#### Statistical analysis

General procedures. Analyses were performed in MATLAB R2026a (MathWorks, Natick, MA, USA). All tests were two-sided. Linear mixed-effects models (LMEs) were fitted in MATLAB using fitlme (Statistics and Machine Learning Toolbox) with restricted maximum likelihood; fixed-effect and planned-contrast denominator degrees of freedom used the Satterthwaite approximation. Categorical fixed effects used effects coding except the pharmacological Treatment factor, for which Before drug was the reference level. The principal session-level LME analyses for Figs. 3d–j and 4 were independently repeated in R 4.6.1 (R Foundation for Statistical Computing, Vienna, Austria) using the lme4, lmerTest, and emmeans packages. The R results closely reproduced the MATLAB estimates and $p$ values, with only negligible numerical differences attributable to independent implementations of the Satterthwaite approximation. Effect estimates and unadjusted 95% confidence intervals (CIs) for key prespecified contrasts are reported in Supplementary Table S6. For LME contrasts, confidence intervals are model-based; for DAT genotype comparisons, Welch-type mean-difference intervals and Hedges' $g$ with 95% confidence intervals are reported. Multiplicity-adjusted inference for prespecified comparison families remains based on Holm-adjusted $p$ values. Exploratory sensitivity analyses incorporating Sex and Sex-related interaction terms into the corresponding statistical models were also performed; the results of these analyses are summarized in Supplementary Table S5. Omnibus fixed-effect $p$ values were unadjusted; $p < 0.05$ was considered statistically significant, whereas omnibus $p$ values between 0.05 and 0.10 were treated as nonsignificant. Prespecified contrasts were corrected with the Holm step-down procedure within the families defined below. Holm-adjusted $p < 0.05$ was considered significant, and $0.05 \le$ Holm-adjusted $p < 0.10$ was interpreted as a trend. Bar graphs show means with upper SEM error bars unless a figure legend states otherwise; spectral curves show geometric means and 95% confidence intervals, and spectral box plots show medians and quartiles. Missing observations were not imputed. The mouse was the statistical unit for behavioral analyses, mouse-level spectral analyses, and DAT expression; complete Before–After session pairs were used for pharmacological comparisons; and individual recording sessions were the observational units for the remaining photometry analyses, as detailed below. Complete statistical results for all analyses corresponding to Figs. 1–5 and Supplementary Figs. S1–S4, including omnibus tests and prespecified comparisons, are provided in Supplementary Table S6.

Figures 1f,g. The two session values within each phase were arithmetically averaged within mouse; mouse was the statistical unit ($N = 10$ per genotype). Repeated-measures models were fitted in MATLAB using fitrm. Phase and Genotype × Phase effects were tested using ranova, whereas the between-subject Genotype effect was obtained using anova. Sphericity was assessed using mauchly, and Greenhouse–Geisser epsilon was obtained using epsilon. Mauchly's test indicated violations of sphericity for both Fig. 1f ($W = 0.0778$, $\chi^2(5) = 42.71$, $p = 4.23 \times 10^{-8}$, $\varepsilon_{GG} = 0.451$) and Fig. 1g ($W = 0.356$, $\chi^2(2) = 17.54$, $p = 1.55 \times 10^{-4}$, $\varepsilon_{GG} = 0.608$); Greenhouse–Geisser corrections were therefore applied to Phase-containing effects. For Fig. 1f, six within-genotype phase comparisons were tested by paired *t* test as a separate six-test Holm family for each genotype; four between-genotype phase-specific comparisons were Welch *t* tests constituting another Holm family. For Fig. 1g, three within-genotype phase comparisons were a separate three-test Holm family for each genotype, and the three phase-specific Welch genotype comparisons formed another family.

Figure 2b. The full time-series spectrum was calculated for each session. Linear session spectra were averaged within MouseID before calculating $\log_{10}$ band power. Control and het-cKO mice were compared by Welch *t* test in each of six bands, separately in the NAc and TS; the six tests within each region formed one Holm family. Mouse was the statistical unit.

Figures 2d–f and S2. Before- and After-drug sessions from the same mouse were sorted by file name, paired by rank, and assigned PairID; only complete pairs were analyzed. For each Drug × Genotype × Region × Metric combination, the session-level LME was Outcome ~ Treatment + (1|MouseID) + (1|PairID), with Before drug as reference and the After-drug coefficient representing After minus Before. For transient amplitude, frequency, and half-width, NAc and TS Treatment contrasts within each Drug × Genotype × Metric combination formed a two-test Holm family. ITI lick frequency had one comparison per Drug × Genotype and therefore required no multiplicity adjustment. When either member of a pair lacked a valid transient amplitude or half-width, that pair was omitted only for the affected metric.

Figures 3d–f. For each Region × Outcome combination, session values were analyzed using USResponse ~ Phase × Genotype + (1|MouseID). Each model contained 80 session observations from 20 mice (20 sessions per Genotype × Phase combination). Four planned contrasts—genotype within Early, genotype within Late, Early versus Late within control, and Early versus Late within het-cKO—formed one Holm family per Region × Outcome.

Figures 3g–j. The pooled dataset comprised session-level observations from Days 8, 9, 11, 12, 15, 17, and 19. Sessions from Day 12 onward were recorded before drug administration, and post-drug sessions were excluded. The Day 12 recording served both as the second Late-phase session and as the pre-drug session; because these designations referred to the same recording session, the Day 12 data were included only once. Thus, each recording session contributed a single observation per region and metric to the pooled analysis. Because the Early- and Late-phase sessions also contributed to the phase-specific analyses in Figs. 3d–f, the datasets underlying Figs. 3d–f and 3g–j partially overlapped and were not treated as independent replications. For each metric, session values were analyzed using MetricValue ~ Region × Genotype + (1|MouseID) + (1|SessionID). SessionID linked the simultaneously recorded NAc and TS observations from the same session. Four planned contrasts—genotype within NAc, genotype within TS, NAc versus TS within control, and NAc versus TS within het-cKO—formed one Holm family per metric.

Figure 4. Separate models were fitted for each Region × preceding Outcome (Air puff or Omission) × Phase combination: CSResponse ~ Sequence × Genotype + (1|MouseID) + (1|SessionID). Sequence had the within-session levels Prior and Subsequent. The two Prior–Subsequent contrasts, one per genotype, formed one Holm family within each panel. The session was the observational unit; $n$ = 20 sessions from $N$ = 10 mice were available in each Genotype × Phase combination.

Figure 5c. The DAT/β-actin ratio from each mouse was the statistical unit (control, $N$ = 6; het-cKO, $N$ = 5). Genotypes were compared within CPu, NAc, TS, and Cb by Welch $t$

tests. The four regional genotype tests formed one Holm family; no between-region contrast was performed.

Figures S1a,b. The primary direct analysis was a Genotype × Region × Band mixed-design repeated-measures ANOVA on mouse-level $\log_{10}$ band power, with Genotype between mice and Region and Band within mice. Greenhouse–Geisser correction was applied to effects involving Band. Within each genotype, NAc–TS differences were tested by paired *t* test within each band; the six bandwise tests formed one Holm family per genotype.

Figures S1c,d. Before- and After-GBR sessions were matched within MouseID by PairID. For the within-genotype analysis, After-minus-Before $\log_{10}$ band-power differences were calculated for each PairID and then averaged within mouse. Each Region × Genotype × Band mean difference was tested against zero by one-sample *t* test; the six bands formed one Holm family within each Region × Genotype. Genotype differences in GBR effects were tested directly in each Region × Band using Log10Power ~ Treatment × Genotype + (1|MouseID) + (1|PairID); the six Treatment × Genotype interaction tests formed one Holm family within each region.

Figures S1e,f. Recovery analyses were restricted to TS and included only mice represented in both the Before-drug and corresponding Recovery conditions defined above. Session $\log_{10}$ band-power values were averaged within MouseID and Condition and paired across conditions. For each Drug × Genotype, six paired bandwise *t* tests formed one Holm family. Genotype differences were tested separately for GBR12909 and SCH23390 by a mixed-design repeated-measures ANOVA with Genotype between mice and Condition and Band within mice; Greenhouse–Geisser correction was applied to Band-containing effects.

Figures S3a–c. NAc and TS observations were paired by SessionID and analyzed separately for Reward, Air puff, and Omission using USResponse ~ Phase × Region × Genotype + (1|MouseID) + (1|SessionID). Each outcome contributed 80 sessions from 20 mice and thus 160 session-by-region observations. The four NAc–TS contrasts within the Genotype × Phase combinations formed one Holm family per outcome.

Figures S4a,b. For each session, outcome-specific CS peaks were pooled by weighting each outcome mean by its trial count, equivalent to averaging all trials regardless of outcome. NAc and TS were modeled separately using PooledPeak ~ Phase × Genotype + (1|MouseID). Four planned contrasts—genotype within Early, genotype within Late,

Early versus Late within control, and Early versus Late within het-cKO—formed one Holm family per region. Each region contributed 80 session observations from 20 mice.

### Supplementary Tables

#### Supplementary Table S1. Genotyping primers

| Primer name | Primer sequence | Amplicon size | PCR condition |
|---|---|---|---|
| ATP2A2 flox allele Forward | 5’-cagaggaccgattgcctctg-3’ | 227 bp for wild-type allele ~327 bp for flox allele | ↓ 95°C, 3 min<br>↓ 95°C, 30 sec<br>↓ 65°C, 30 sec } x 35 cycles<br>↓ 72°C, 30 sec<br>↓ 72°C, 5 min<br>15°C hold |
| ATP2A2 flox allele Reverse | 5’--ccattacagatggttgtaagtcacc~ 3’ | | |
| Cre transgene Forward | 5’-acctgatggacatgttcagggatcg-3’ | 107 bp | ↓ 95°C, 3min<br>↓ 95°C, 30sec<br>↓ 60°C, 30sec } x 35 cycles<br>↓ 72°C, 30sec<br>↓ 72°C, 2min<br>15°C hold |
| Cre transgene Reverse | 5’- tccggttattcaacttgcaccatgc-3’ | | |

#### Supplementary Table S2. Age and sex composition of mice used for the analyses in Figs. 3 and 4

| Genotype | N | Age at first Early-phase recording (weeks), mean ± SD | Range (weeks) | Sex composition |
|---|---|---|---|---|
| Control | 10 | 37.4 ± 9.3 | 23.0–50.4 | 6 females, 4 males |
| het-cKO | 10 | 31.2 ± 7.7 | 16.6–43.4 | 5 females, 5 males |

Note: Data are presented as mean ± SD. Age was calculated at the first Early-phase recording session for each mouse. The mean age did not differ significantly between genotypes (Welch *t* test, $t(17.39) = 1.63$, $p = 0.121$).

#### Supplementary Table S3. Experimental design of the classical conditioning task

| Parameter | Pre-training | Training | Test |
|---|---|---|---|
| CS duration (s) | 3 | 6 | 10 |
| Delay period (s) | — | 3 | 3 |
| Response period (s) | 3 | 3 | 7 |
| ITI (s) | 3 | 6 | 20–35 |
| Trials/session | 200 | 150 | 50 |
| Reinforcement contingency | First lick during the CS triggered reward | First response-period lick triggered reward | First response-period lick triggered reward (80%), air puff (10%), or omission (10%) |

Test outcomes were presented in pseudorandom order while preserving the indicated session-level frequencies.

#### Supplementary Table S4. Experimental-day assignments for phase-based, pooled, pharmacological, and recovery analyses

| Day | Session designation | Drug administered after session | Phase-based analyses | Fig. 3g–j pooled dataset | Fig. S1e,f recovery role |
|---|---|---|---|---|---|
| 8 | Early 1 | — | Early | Included | — |
| 9 | Early 2 | — | Early | Included | — |
| 10 | Intervening Test session | — | Not included | Not included | — |
| 11 | Late 1 | — | Late | Included | — |
| 12 | Late 2 / Before GBR | GBR12909 | Late | Included once | — |
| 15 | Before SCH | SCH23390 | — | Included | Recovery after preceding GBR12909 |
| 16 | Drug-free Test session | — | — | Not included | Not used for Figs. S1e,f |
| 17 | Before GBR | GBR12909 | — | Included | Recovery after preceding SCH23390 |
| 18 | Drug-free Test session | — | — | Not included | Not used for Figs. S1e,f |
| 19 | Before SCH | SCH23390 | — | Included | Recovery after preceding GBR12909 |

Note: The day 12 recording was a single session that served both as the second Late-phase session and as the Before-drug baseline for the first GBR12909 administration; it was therefore counted only once in the Fig. 3g–j pooled dataset. Recovery analyses in Figs. S1e,f used the indicated later pre-drug recordings rather than the separate drug-free Test sessions on days 16 and 18.

#### Supplementary Table S5. Exploratory sex-sensitivity analyses

| Figure(s) | Analysis | Sex-sensitivity model / correction | Summary |
|---|---|---|---|
| Figs. 2d–f and S2 | Pharmacological Before–After analyses | Exploratory sex sensitivity was performed on mouse-level treatment effects derived from complete Before–After pairs. Sex-related analyses were secondary and did not determine figure annotations. | Exploratory sensitivity analysis; primary inference remained the PairID-aware session-level LME. |
| Figs. 3d–f | Outcome-evoked response peaks | USResponse ~ Phase × Genotype × Sex + (1\|MouseID). For each Sex-containing term, Holm correction was applied across the six Region × Outcome panels. | Genotype × Sex and Phase × Genotype × Sex terms were not significant after correction; the genotype-related conclusions of the primary analysis were unchanged. |
| Figs. 3g–j | Reward-response and ITI-transient metrics | MetricValue ~ Region × Genotype × Sex + (1\|MouseID) + (1\|SessionID). For each Sex-containing term, Holm correction was applied across the four metrics. | Genotype × Sex and Region × Genotype × Sex terms were not significant after correction. Some Region × Sex effects were observed in session-level analyses but were not reproduced consistently in mouse-level sensitivity analyses and did not indicate sex-dependent genotype effects. |
| Fig. 4 | Trial-history-dependent CS responses | CSResponse ~ Sequence × Genotype × Sex + (1\|MouseID) + (1\|SessionID), evaluated separately for each Region × preceding outcome × Phase combination; Sex-containing terms were treated as exploratory. | No sex-related result altered the interpretation of the primary Sequence × Genotype analyses; primary inference was therefore based on the prespecified sex-collapsed models. |
| Figs. S3a–c | Direct regional comparison of outcome responses | USResponse ~ Phase × Region × Genotype × Sex + random intercepts corresponding to the primary session-level analysis; Sex-containing terms were exploratory. | No sex-dependent modification was used to support the reported regional or genotype effects. |
| Figs. S4a,b | Outcome-pooled CS responses | PooledPeak ~ Phase × Genotype × Sex + (1\|MouseID). Sex-containing terms were exploratory. | Primary conclusions were based on the prespecified Phase × Genotype model; the sensitivity analysis did not provide evidence requiring a sex-specific reinterpretation. |

Sex-sensitivity analyses were exploratory and were not used to determine the significance annotations in the figures. They were intended to test whether inclusion of Sex materially altered genotype-related conclusions rather than to provide adequately powered sex-stratified inference.

#### Supplementary Table S6. Statistical results for Figs. 1–5 and Supplementary Figs. S1–S4

| Figure | Analysis / metric | Effect or planned comparison | Statistic / effect estimate (95% CI) | *p* value | Interpretation |
|---|---|---|---|---|---|
| Fig. 1f | Response-period Δlick frequency | Phase omnibus (GG-corrected) | $F(1.353, 24.358) = 27.29$ | $p_{GG} = 5.04 \times 10^{-6}$ | Significant |
| | | Genotype omnibus | $F(1, 18) = 1.21$ | $p = 0.286$ | n.s. |
| | | Genotype × Phase omnibus (GG-corrected) | $F(1.353, 24.358) = 0.84$ | $p_{GG} = 0.402$ | n.s. |
| | Sphericity | Mauchly test | $W = 0.0778$; $\chi^2(5) = 42.71$ | $p = 4.23 \times 10^{-8}$; $\varepsilon_{GG} = 0.451$ | GG correction applied |
| | Within-genotype planned phase contrasts | Control | paired *t* tests | 5 of 6 contrasts $p_{Holm} = 0.00105–0.0186$ | Training–Test Early n.s. |
| | | het-cKO | paired *t* tests | $p_{Holm} = 0.0164–0.0436$ for 5 significant contrasts; Training–Test Early, $p_{Holm} = 0.0888$ | 5 of 6 contrasts significant; Training–Test Early showed a trend |
| Fig. 1g | Anticipatory Δlick frequency | Phase omnibus (GG-corrected) | $F(1.217, 21.902) = 23.09$ | $p_{GG} = 3.82 \times 10^{-5}$ | Significant |
| | | Genotype omnibus | $F(1, 18) = 0.0368$ | $p = 0.850$ | n.s. |
| | | Genotype × Phase omnibus (GG-corrected) | $F(1.217, 21.902) = 0.0167$ | $p_{GG} = 0.932$ | n.s. |
| | Sphericity | Mauchly test | $W = 0.356$; $\chi^2(2) = 17.54$ | $p = 1.55 \times 10^{-4}$; $\varepsilon_{GG} = 0.608$ | GG correction applied |
| | Planned phase contrasts, pooled across genotype | Training vs Test Early | $t(19) = -4.53$ | $p_{Holm} = 4.55 \times 10^{-4}$ | Significant |
| | | Training vs Test Late | $t(19) = -5.39$ | $p_{Holm} = 1.01 \times 10^{-4}$ | Significant |
| | | Test Early vs Test Late | $t(19) = -4.31$ | $p_{Holm} = 4.55 \times 10^{-4}$ | Significant |
| Fig. 2b | Mouse-level $\log_{10}$ band power | Genotype comparisons, NAc (six bands) | Welch *t* tests | all $p_{Holm} \geq 0.200$ | No significant difference or trend |
| | | Genotype comparisons, TS (six bands) | Welch *t* tests | all $p_{Holm} \geq 0.838$ | No significant difference or trend |
| Fig. 2d | NAc transient frequency, GBR | Treatment, control | $F(1, 26.45) = 13.72$ | $p_{Holm} = 0.00197$ | Increased after GBR |
| | | Treatment, het-cKO | $F(1, 20.04) = 7.93$ | $p_{Holm} = 0.0155$ | Increased after GBR |
| Fig. 2d | NAc transient frequency, SCH | Treatment, control | $F(1, 24.19) = 34.12$ | $p_{Holm} = 4.90 \times 10^{-6}$ | Decreased after SCH |
| | | Treatment, het-cKO | $F(1, 18.07) = 14.21$ | $p_{Holm} = 0.00139$ | Decreased after SCH |
| Fig. 2e | NAc transient half-width, GBR | Treatment, control | $F(1, 25.18) = 23.90$ | $p_{Holm} = 4.88 \times 10^{-5}$ | Increased after GBR |
| | | Treatment, het-cKO | $F(1, 22.31) = 18.75$ | $p_{Holm} = 4.34 \times 10^{-4}$ | Increased after GBR |
| | NAc transient half-width, SCH | Treatment, control | LME treatment contrast | $p > 0.10$ | n.s. |
| | | Treatment, het-cKO | $F(1, 13.00) = 6.72$ | $p_{Holm} = 0.0446$ | Decreased after SCH |
| Fig. 2f | ITI lick frequency, GBR | Treatment, control | $F(1, 32.00) = 11.28$ | $p = 0.00204$ | Increased after GBR |
| | | Treatment, het-cKO | $F(1, 15.00) = 20.23$ | $p = 4.25 \times 10^{-4}$ | Increased after GBR |
| | ITI lick frequency, SCH | Treatment, control | $F(1, 15.00) = 66.41$ | $p = 6.85 \times 10^{-7}$ | Decreased after SCH |
| | | Treatment, het-cKO | $F(1, 19.47) = 29.82$ | $p = 2.64 \times 10^{-5}$ | Decreased after SCH |
| Fig. S1a,b | Regional spectral organization, control | Region × Band | $F(5, 45) = 11.29$ | $p_{GG} = 0.00217$ | Significant |
| | Regional spectral organization, het-cKO | Region × Band | $F(5, 45) = 15.18$ | $p_{GG} = 7.66 \times 10^{-5}$ | Significant |
| | Direct Genotype × Region × Band analysis | Genotype | $F(1, 18) = 4.16$ | $p = 0.0564$ | n.s. |

| Figure | Analysis / metric | Effect or planned comparison | Statistic / effect estimate (95% CI) | *p* value | Interpretation |
|---|---|---|---|---|---|
| | | Genotype × Region | $F(1, 18) = 3.30$ | $p = 0.0858$ | n.s. |
| | | Genotype × Band | $F(5, 90) = 2.72$ | $p_{GG} = 0.0803$ | n.s. |
| | | Genotype × Region × Band | $F(5, 90) = 1.51$ | $p_{GG} = 0.235$ | n.s. |
| Fig. S1c,d | GBR spectral sensitivity | Treatment × Genotype, all bands/regions | Bandwise models | all $p_{Holm} \geq 0.243$ | No genotype difference in GBR sensitivity |
| Fig. S1e,f | Recovery vs Before drug | Bandwise Before–Recovery contrasts | Paired tests | all $p_{Holm} \geq 0.634$ | No residual band-power difference or trend |
| Fig. S1e,f | Recovery analysis | Genotype × Condition and Genotype × Condition × Band | Mixed repeated-measures ANOVA | all $p > 0.05$ | n.s. |
| Fig. S2 | Transient amplitude | All Drug × Genotype × Region treatment contrasts | PairID-aware LME | all $p_{Holm} \geq 0.204$ | No drug effect on amplitude |
| | TS transient frequency, GBR | Treatment, control | PairID-aware LME | $p_{Holm} \geq 0.10$ | n.s. |
| | | Treatment, het-cKO | $F(1, 20.15) = 8.74$ | $p_{Holm} = 0.0155$ | Increased after GBR |
| | TS transient frequency, SCH | Treatment, control | $F(1, 15.00) = 89.36$ | $p_{Holm} = 2.08 \times 10^{-7}$ | Decreased after SCH |
| | | Treatment, het-cKO | $F(1, 13.00) = 25.27$ | $p_{Holm} = 4.63 \times 10^{-4}$ | Decreased after SCH |
| | TS transient half-width, GBR | Treatment, control and het-cKO | PairID-aware LME | both $p_{Holm} < 0.05$ | Increased after GBR |
| | TS transient half-width, SCH | Treatment, control and het-cKO | PairID-aware LME | both $p_{Holm} \geq 0.10$ | n.s. |
| Fig. 3d | Reward peak, NAc | Genotype | $F(1, 18.00) = 4.20$ | $p = 0.0553$ | n.s. |
| | | Phase × Genotype | LME omnibus | $p = 0.648$ | n.s. |
| | | Early: control vs het-cKO | planned contrast; raw $p = 0.0488$ | $p_{Holm} = 0.195$ | n.s. after correction |
| | Reward peak, TS | Phase, Genotype, interaction and planned contrasts | LME / Holm contrasts | all $p \geq 0.10$ after correction | n.s. |
| Fig. 3e | Air-puff peak, NAc | Phase | $F(1, 58.00) = 35.03$ | $p = 1.86 \times 10^{-7}$ | Significant |
| | Air-puff peak, TS | Phase | $F(1, 58.00) = 43.50$ | $p = 1.39 \times 10^{-8}$ | Significant |
| | Air-puff peak, NAc | Phase × Genotype | $F(1, 58.00) = 2.99$ | $p = 0.0890$ | n.s. |
| | | Early: control vs het-cKO | planned contrast | $p_{Holm} = 0.0650$ | Trend |
| | | Early vs Late, control | $t(58.00) = 2.96$ | $p_{Holm} = 0.0133$ | Early > Late |
| | | Early vs Late, het-cKO | $t(58.00) = 5.41$ | $p_{Holm} = 5.03 \times 10^{-6}$ | Early > Late |
| | Air-puff peak, TS | Early vs Late, control | $t(58.00) = 5.34$ | $p_{Holm} = 6.49 \times 10^{-6}$ | Early > Late |
| | | Early vs Late, het-cKO | $t(58.00) = 3.99$ | $p_{Holm} = 5.67 \times 10^{-4}$ | Early > Late |
| Fig. 3f | Omission peak, NAc | Genotype | $F(1, 18.00) = 4.94$ | $p = 0.0393$ | Significant |
| | | Phase × Genotype | $F(1, 58.00) = 4.59$ | $p = 0.0365$ | Significant |
| | | Early: control vs het-cKO | $t(45.88) = -3.08$ | $p_{Holm} = 0.0140$ | het-cKO > control |
| | | Early vs Late, het-cKO | $t(58.00) = 2.69$ | $p_{Holm} = 0.0277$ | Early > Late |
| Fig. 3f | Omission peak, TS | Phase | $F(1, 58.00) = 9.52$ | $p = 0.00312$ | Significant |
| | | Genotype | $F(1, 18.00) = 6.67$ | $p = 0.0188$ | Significant |
| | | Phase × Genotype | LME omnibus | $p = 0.129$ | n.s. |
| | | Early: control vs het-cKO | $t(44.97) = -2.97$ | $p_{Holm} = 0.0143$ | het-cKO > control |
| | | Early vs Late, het-cKO | $t(58.00) = 3.27$ | $p_{Holm} = 0.00724$ | Early > Late |
| Fig. 3g | Reward-response amplitude | Region | $F(1, 252.92) = 25.09$ | $p = 1.03 \times 10^{-6}$ | Significant |
| | | Genotype | $F(1, 17.82) = 4.46$ | $p = 0.0491$ | Significant |

| Figure | Analysis / metric | Effect or planned comparison | Statistic / effect estimate (95% CI) | *p* value | Interpretation |
|---|---|---|---|---|---|
| | | NAc: control vs het-cKO | $t(22.71) = -2.43$<br>Δ(het-cKO − control) = 0.687 z-score (95% CI 0.102–1.273) | $p_{Holm} = 0.0469$ | het-cKO > control |
| | | NAc vs TS, control | $t(252.94) = 2.63$ | $p_{Holm} = 0.0273$ | NAc > TS |
| | | NAc vs TS, het-cKO | $t(252.90) = 4.44$ | $p_{Holm} = 5.27 \times 10^{-5}$ | NAc > TS |
| Fig. 3h | Reward-response half-width | Region | $F(1, 252.64) = 136.04$ | $p = 1.96 \times 10^{-25}$ | Significant |
| | | NAc vs TS, control | $t(252.66) = 9.01$ | $p_{Holm} = 2.10 \times 10^{-16}$ | NAc > TS |
| | | NAc vs TS, het-cKO | $t(252.62) = 7.49$ | $p_{Holm} = 3.36 \times 10^{-12}$ | NAc > TS |
| | | Genotype and Region × Genotype | LME omnibus | both $p > 0.05$ | n.s. |
| Fig. 3i | ITI transient frequency | Region | $F(1, 253.74) = 79.27$ | $p = 1.06 \times 10^{-16}$ | Significant |
| | | Genotype | $F(1, 17.63) = 3.45$ | $p = 0.0799$ | n.s. |
| | | NAc vs TS, control | $t(253.74) = -7.05$ | $p_{Holm} = 6.83 \times 10^{-11}$ | TS > NAc |
| | | NAc vs TS, het-cKO | $t(253.74) = -5.55$ | $p_{Holm} = 2.11 \times 10^{-7}$ | TS > NAc |
| | | NAc: control vs het-cKO | $t(23.4) = -2.09$<br>Δ(het-cKO − control) = 0.789 events/min (95% CI 0.009–1.569) | $p_{Holm} = 0.0950$ | Trend; het-cKO > control |
| Fig. 3j | ITI transient half-width | Region | $F(1, 245.51) = 80.50$ | $p = 7.70 \times 10^{-17}$ | Significant |
| | | Genotype | $F(1, 16.32) = 4.37$ | $p = 0.0527$ | n.s. |
| | | NAc vs TS, control | $t(249.02) = 5.43$ | $p_{Holm} = 4.00 \times 10^{-7}$ | NAc > TS |
| | | NAc vs TS, het-cKO | $t(241.24) = 7.29$ | $p_{Holm} = 1.72 \times 10^{-11}$ | NAc > TS |
| | | NAc: control vs het-cKO | $t(39.2) = -2.31$<br>Δ(het-cKO − control) = 0.0421 s (95% CI 0.0053–0.0788) | $p_{Holm} = 0.0520$ | Trend; het-cKO > control |
| Fig. S3a | Reward response, direct regional model | Region | $F(1, 134) = 7.10$ | $p = 0.00868$ | Significant |
| | | Genotype | $F(1, 18) = 5.11$ | $p = 0.0364$ | Significant |
| | | Region × Genotype | $F(1, 134) = 3.19$ | $p = 0.0765$ | n.s. |
| | Reward response | NAc vs TS, het-cKO Early and Late | planned contrasts | both $p_{Holm} = 0.0978$ | Trend |
| Fig. S3b | Air-puff response, direct regional model | Phase | $F(1, 134) = 39.83$ | $p = 3.76 \times 10^{-9}$ | Significant |
| | | Region | $F(1, 134) = 62.00$ | $p = 1.04 \times 10^{-12}$ | Significant |
| | | Region × Genotype | $F(1, 134) = 6.86$ | $p = 0.00984$ | Significant |
| | Air-puff response | NAc vs TS, all Genotype × Phase cells | planned contrasts | all $p_{Holm} \leq 0.0167$ | TS > NAc |
| Fig. S3c | Omission response, direct regional model | Phase | $F(1, 58) = 7.03$ | $p = 0.0103$ | Significant |
| | | Region | $F(1, 76) = 29.25$ | $p = 7.10 \times 10^{-7}$ | Significant |
| | | Genotype | $F(1, 18) = 8.86$ | $p = 0.00808$ | Significant |
| | | Phase × Genotype | $F(1, 58) = 4.78$ | $p = 0.0328$ | Significant |
| | Omission response | NAc vs TS, all Genotype × Phase cells | planned contrasts | $p_{Holm} = 0.00200–0.0460$ | TS > NAc |
| Fig. S4a | Pooled CS-response peak, NAc | Phase | $F(1, 58) = 17.03$ | $p = 1.19 \times 10^{-4}$ | Significant |
| | | Genotype | LME omnibus | $p = 0.340$ | n.s. |

| Figure | Analysis / metric | Effect or planned comparison | Statistic / effect estimate (95% CI) | *p* value | Interpretation |
|---|---|---|---|---|---|
| | | Phase × Genotype | LME omnibus | $p = 0.662$ | n.s. |
| | | Early vs Late, control | planned contrast | $p_{Holm} = 0.00820$ | Late > Early |
| | | Early vs Late, het-cKO | planned contrast | $p_{Holm} = 0.0347$ | Late > Early |
| Fig. S4b | Pooled CS-response peak, TS | Phase / Genotype / interaction | LME omnibus | $p$ = 0.698 / 0.882 / 0.834 | All n.s. |
| Fig. S4b | Pooled CS-response peak, TS | All four planned contrasts | Holm contrasts | all $p_{Holm} = 1$ | n.s. |
| Fig. 4a | NAc Air puff→Reward, Early | Sequence | $F(1, 38.00) = 13.25$ | $p = 8.07 \times 10^{-4}$ | Significant |
| | | Genotype | $F(1, 18.00) = 5.43$ | $p = 0.0316$ | Significant |
| | | Prior vs Subsequent, control | $t(38.00) = 2.59$<br>Δ(Prior − Subsequent) = 0.496 z-score (95% CI 0.109–0.883) | $p_{Holm} = 0.0269$ | Prior > Subsequent |
| | | Prior vs Subsequent, het-cKO | $t(38.00) = 2.56$<br>Δ(Prior − Subsequent) = 0.489 z-score (95% CI 0.102–0.876) | $p_{Holm} = 0.0269$ | Prior > Subsequent |
| | NAc Air puff→Reward, Late | Sequence | $F(1, 38.00) = 7.70$ | $p = 0.00851$ | Significant |
| | | Prior vs Subsequent, control | $t(38.00) = 0.974$<br>Δ(Prior − Subsequent) = 0.157 z-score (95% CI −0.169–0.483) | $p_{Holm} = 0.336$ | n.s. |
| | | Prior vs Subsequent, het-cKO | $t(38.00) = 2.95$<br>Δ(Prior − Subsequent) = 0.476 z-score (95% CI 0.149–0.802) | $p_{Holm} = 0.0108$ | Prior > Subsequent |
| | | Sequence × Genotype | $F(1, 38.00) = 1.95$ | $p = 0.170$ | n.s. |
| Fig. 4b | NAc Omission→Reward, Early | Genotype | $F(1, 18.00) = 3.87$ | $p = 0.0647$ | n.s. |
| | NAc Omission→Reward, remaining analyses | Sequence, interaction and planned contrasts | LME / Holm contrasts | all $p \geq 0.10$ after correction | n.s. |
| Fig. 4c | TS Air puff→Reward, Early | Sequence × Genotype | $F(1, 38.00) = 3.42$ | $p = 0.0722$ | n.s. |
| | TS Air puff→Reward, remaining analyses | Sequence, genotype, interaction and planned contrasts | LME / Holm contrasts | all $p \geq 0.10$ after correction | n.s. |
| Fig. 4d | TS Omission→Reward, Early | Sequence × Genotype | $F(1, 38.00) = 2.88$ | $p = 0.0980$ | n.s. |
| | TS Omission→Reward, remaining analyses | Sequence, genotype, interaction and planned contrasts | LME / Holm contrasts | all $p \geq 0.10$ after correction | n.s. |
| Fig. 4a–d | All eight Region × preceding outcome × Phase models | Sequence × Genotype | LME omnibus | all $p > 0.05$ | No significant interaction |
| Fig. 5c | DAT/β-actin, CPu | Control vs het-cKO | Welch $t(9.79) = 1.11$ Δ(het-cKO − control) = −0.179 (95% CI [−0.541, 0.182]); Hedges' $g$ = −0.59 (95% CI [−1.65, 0.50]) | $p_{Holm} = 1$ | n.s. |
| | DAT/β-actin, NAc | Control vs het-cKO | Welch $t(7.73) = 1.01$ Δ(het-cKO − control) = −0.0867 (95% CI [−0.286, 0.113]); Hedges' $g$ = −0.53 (95% CI [−1.62, 0.60]) | $p_{Holm} = 1$ | n.s. |
| | DAT/β-actin, TS | Control vs het-cKO | Welch $t(9.84) = 0.842$ Δ(het-cKO − control) = −0.0508 (95% CI [−0.186, 0.0840]); Hedges' $g$ = −0.45 (95% CI [−1.50, 0.62]) | $p_{Holm} = 1$ | n.s. |
| | DAT/β-actin, Cb | Control vs het-cKO | Welch $t(6.30) = -0.697$ Δ(het-cKO − control) = 0.0114 (95% CI [−0.0282, 0.0510]); Hedges' $g$ = 0.37 (95% CI [−0.69, 1.42]) | $p_{Holm} = 1$ | n.s. |

Notes: Omnibus fixed-effect *p* values were unadjusted and values between 0.05 and 0.10 were treated as nonsignificant. $p_{Holm}$ denotes Holm-adjusted *p* values for the prespecified comparison family defined in the Statistical analysis section. $p_{GG}$ denotes Greenhouse–Geisser-corrected *p* values. The term "trend" was reserved for prespecified contrasts with $0.05 \leq p_{Holm} < 0.10$. "n.s." indicates not significant. Where multiple nonsignificant tests are summarized in a single row, the range/inequality shown reflects the final audited analysis rather than an invented point estimate. LME = linear mixed-effects model; NAc = nucleus accumbens; TS = tail of the striatum; Cb = cerebellum. Effect estimates are reported in the direction specified in the Statistic column. Confidence intervals are unadjusted 95% confidence intervals (model-based for LME contrasts and Welch-type for DAT mean differences); Holm-adjusted *p* values remain the basis for inference. For DAT comparisons, Hedges' *g* uses the pooled standard deviation with the small-sample correction, and its 95% confidence interval was obtained by noncentral-t inversion.

#### Supplementary Figure Legends

##### Figure S1. Regional spectral organization, GBR12909-induced spectral changes, and recovery.

Power spectra and band power in six frequency bands. Spectral traces show means and 95% confidence intervals. Box plots show the median, interquartile range, and minimum-to-maximum whiskers. (a, b) Power spectra and band power in the NAc and TS in control (a; $N$ = 10 mice) and het-cKO mice (b; $N$ = 10 mice). Mouse-level mean spectra were entered into Region × Band repeated-measures ANOVAs and paired $t$ tests; six bandwise comparisons were Holm-adjusted within each genotype. Region × Band interactions were significant in controls ($F(5, 45) = 11.29$, $p_{GG} = 0.00217$; Greenhouse–Geisser correction) and het-cKO mice ($F(5, 45) = 15.18$, $p_{GG} = 7.66 \times 10^{-5}$). In controls, NAc power exceeded TS power at 0.01–0.03 Hz ($p_{Holm} = 0.0313$), whereas TS power exceeded NAc power at 0.3–1.0 and 1.0–3.0 Hz ($p_{Holm} = 0.0291$ and $1.25 \times 10^{-4}$). In het-cKO mice, NAc power was greater at 0.01–0.03 and 0.03–0.1 Hz ($p_{Holm} = 0.00720$ and $6.59 \times 10^{-4}$), whereas TS power was greater at 1.0–3.0 Hz ($p_{Holm} = 0.00468$); regional differences at 0.3–1.0 and 3.0–10 Hz showed trends (both $p_{Holm} = 0.0640$). A direct mouse-level Genotype × Region × Band analysis showed trends for Genotype ($F(1, 18) = 4.16$, $p = 0.0564$), Genotype × Region ($F(1, 18) = 3.30$, $p = 0.0858$), and Genotype × Band ($F(5, 90) = 2.72$, $p_{GG} = 0.0803$), but not for Genotype × Region × Band ($F(5, 90) = 1.51$, $p_{GG} = 0.235$). (c, d) Power spectra and band power in six frequency bands before and after GBR12909 administration in the NAc and TS of control (c) and het-cKO mice (d). Within each PairID, After-minus-Before differences were calculated, averaged within mouse, and tested against zero; six bandwise comparisons were Holm-adjusted within each Region × Genotype combination (control, $n$ = 17 matched pairs from $N$ = 7 mice; het-cKO, $n$ = 16 matched pairs from $N$ = 9 mice). In control mice, NAc power showed an increasing trend at 0.03–0.1 Hz ($p_{Holm} = 0.0915$), whereas TS power increased at 0.03–0.1 and 0.1–0.3 Hz ($p_{Holm} = 0.00179$ and $0.0163$) and decreased at 1.0–3.0 and 3.0–10 Hz ($p_{Holm} = 6.14 \times 10^{-4}$ and $0.00179$). In het-cKO mice, NAc power decreased at 1.0–3.0 Hz ($p_{Holm} = 0.0335$); TS power decreased at 3.0–10 Hz ($p_{Holm} = 0.00242$) and showed a decreasing trend at 1.0–3.0 Hz ($p_{Holm} = 0.0918$). Treatment × Genotype interactions were nonsignificant in every band in both regions (all $p_{Holm} \geq 0.243$). (e, f) Power spectra and band power in six frequency bands in the TS under the Before-drug and Recovery conditions after GBR12909 (upper) and SCH23390 (lower), shown for control (e; GBR, $N = 7$ mice; SCH, $N = 7$ mice) and het-cKO mice (f; GBR, $N = 8$ mice; SCH, $N = 5$ mice). Mouse-level paired tests showed no bandwise difference or trend (all $p_{\mathrm{Holm}} \geq 0.634$). For GBR12909, Genotype × Condition ($F(1, 13) = 1.91$, $p = 0.191$) and Genotype × Condition × Band ($F(5, 65) = 1.60$, $p_{\mathrm{GG}} = 0.222$) were nonsignificant. For SCH23390, Genotype showed a trend ($F(1, 10) = 3.71$, $p = 0.0830$), whereas Genotype × Condition ($F(1, 10) = 1.20$, $p = 0.299$) and Genotype × Condition × Band ($F(5, 50) = 0.540$, $p_{\mathrm{GG}} = 0.545$) were nonsignificant. *$p < 0.05$; **$p < 0.01$; ***$p < 0.001$; †$0.05 \leq p < 0.1$ (Holm-adjusted).

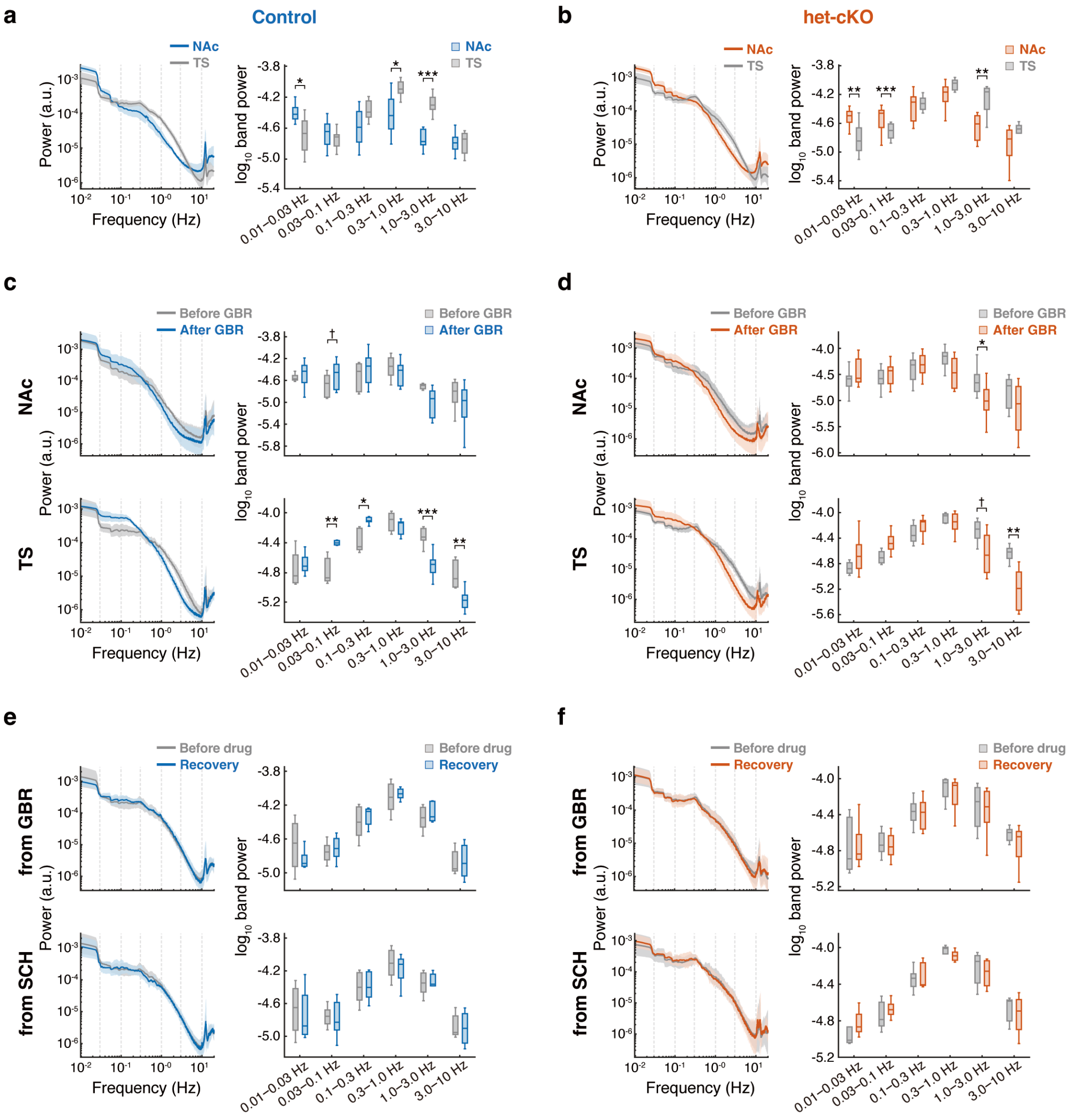

**Figure S2. Pharmacological effects on spontaneous dopamine-transient amplitude and TS transient dynamics.**

Bars show means + SEM across matched recording sessions for GBR12909 (top row) and SCH23390 (bottom row): (a) NAc ITI transient amplitude; (b) TS ITI transient amplitude; (c) TS ITI transient frequency; and (d) TS ITI transient half-width. Within each panel, control and het-cKO groups are shown side by side, with Before and After bars for each genotype. Complete matched Before–After session pairs were analyzed using the linear mixed-effects model Outcome ~ Treatment + (1|MouseID) + (1|PairID), fitted by restricted maximum likelihood with Satterthwaite degrees of freedom. Treatment contrasts for the NAc and TS were Holm-adjusted as a two-test family within each Drug × Genotype × Metric combination. ITI transient amplitude did not change significantly in either region after either drug (all $p_{\text{Holm}} \geq 0.204$). TS transient frequency increased after GBR12909 in het-cKO mice ($F(1, 20.15) = 8.74$, $p_{\text{Holm}} = 0.0155$), but not in controls ($p_{\text{Holm}} = 0.148$), and decreased after SCH23390 in control ($F(1, 15.00) = 89.36$, $p_{\text{Holm}} = 2.08 \times 10^{-7}$) and het-cKO mice ($F(1, 13.00) = 25.27$, $p_{\text{Holm}} = 4.63 \times 10^{-4}$). TS transient half-width increased after GBR12909 in control ($F(1, 27.23) = 38.26$, $p_{\text{Holm}} = 2.50 \times 10^{-6}$) and het-cKO mice ($F(1, 25.40) = 18.56$, $p_{\text{Holm}} = 4.34 \times 10^{-4}$), but did not change after SCH23390 (control, $p_{\text{Holm}} = 0.313$; het-cKO, $p_{\text{Holm}} = 0.165$). Maximum available sample sizes were $n = 17$ matched pairs from $N = 7$ mice (control, GBR), $n = 16$ matched pairs from $N = 9$ mice (het-cKO, GBR), $n = 16$ matched pairs from $N = 7$ mice (control, SCH), and $n = 14$ matched pairs from $N = 8$ mice (het-cKO, SCH). Sample sizes for transient amplitude and half-width were smaller when either recording in a matched Before–After pair contained no valid transient from which the measure could be calculated. *$p < 0.05$; ***$p < 0.001$ (Holm-adjusted).

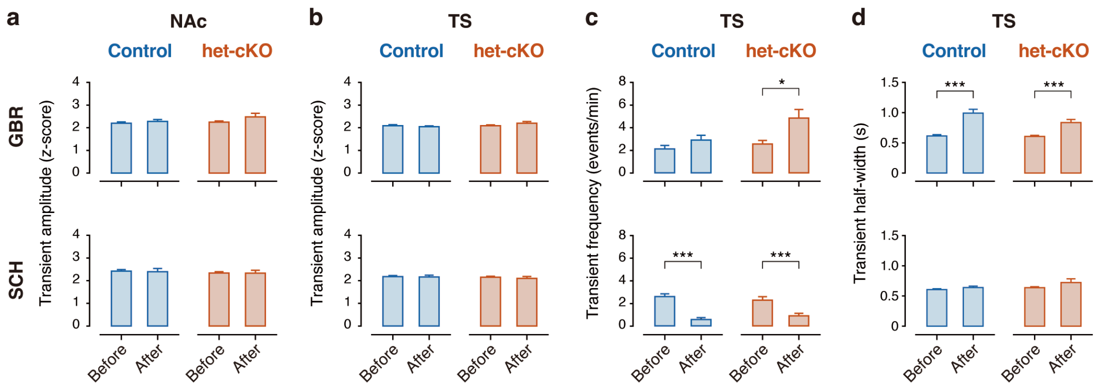

**Figure S3. Direct regional comparison of outcome-evoked dopamine responses.** NAc and TS US-response amplitudes are shown for Reward (a), Air puff (b), and Omission trials (c) during the Early and Late phases, separately for control (top) and het-cKO mice (bottom). For each outcome, 80 recording sessions from 20 mice ($N = 10$ mice per genotype) yielded 160 session-by-region observations because the NAc and TS were recorded simultaneously in each session. Session-level data were analyzed using the linear mixed-effects model USResponse ~ Phase × Region × Genotype + (1|MouseID) + (1|SessionID); NAc–TS contrasts within each Genotype × Phase combination were Holm-adjusted across four comparisons per outcome. Reward responses showed Region ($F(1, 134) = 7.10$, $p = 0.00868$) and Genotype effects ($F(1, 18) = 5.11$, $p = 0.0364$), with a Region × Genotype trend ($F(1, 134) = 3.19$, $p = 0.0765$). NAc responses tended to exceed TS responses in het-cKO mice during both Early and Late (both $p_{Holm} = 0.0978$). Air-puff responses showed Phase ($F(1, 134) = 39.83$, $p = 3.76 \times 10^{-9}$), Region ($F(1, 134) = 62.00$, $p = 1.04 \times 10^{-12}$), and Region × Genotype effects ($F(1, 134) = 6.86$, $p = 0.00984$). TS responses exceeded NAc responses in all four Genotype × Phase combinations (control Early, $p_{Holm} = 6.15 \times 10^{-9}$; control Late, $p_{Holm} = 3.08 \times 10^{-4}$; het-cKO Early and Late, both $p_{Holm} = 0.0167$). Omission responses showed Phase ($F(1, 58) = 7.03$, $p = 0.0103$), Region ($F(1, 76) = 29.25$, $p = 7.10 \times 10^{-7}$), Genotype ($F(1, 18) = 8.86$, $p = 0.00808$), and Phase × Genotype effects ($F(1, 58) = 4.78$, $p = 0.0328$). TS responses exceeded NAc responses in all four Genotype × Phase combinations (control Early, $p_{Holm} = 0.00200$; control Late, $p_{Holm} = 0.0460$; het-cKO Early, $p_{Holm} = 0.0347$; het-cKO Late, $p_{Holm} = 0.0243$). Bars show means + SEM across sessions. *$p < 0.05$; **$p < 0.01$; ***$p < 0.001$; †$0.05 \leq p < 0.1$ (Holm-adjusted).

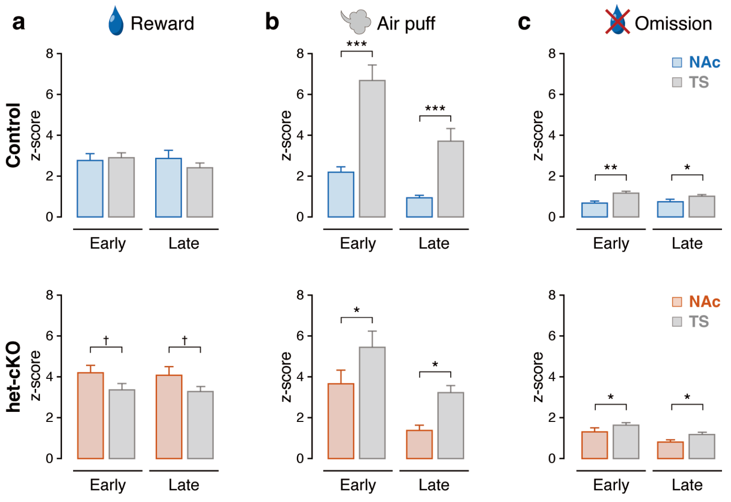

**Figure S4. Phase-dependent conditioned-stimulus responses pooled across outcomes.**

CS-response peaks were calculated from stimulus-onset-aligned dopamine signals averaged across all trials, irrespective of outcome, and compared between the Early and Late phases in the NAc (a) and TS (b). Each region included 80 session-level observations from 20 mice ($N$ = 10 mice per genotype). Session-level data were analyzed using the linear mixed-effects model CSResponse ~ Phase × Genotype + (1|MouseID), with four planned comparisons Holm-adjusted within each region. In the NAc, the Phase effect was significant ($F(1, 58) = 17.03$, $p = 1.19 \times 10^{-4}$), whereas Genotype ($F(1, 18) = 0.959$, $p = 0.340$) and Phase × Genotype ($F(1, 58) = 0.193$, $p = 0.662$) were not. CS responses increased from Early to Late in control ($p_{Holm} = 0.00820$) and het-cKO mice ($p_{Holm} = 0.0347$), with no genotype difference within either phase. In the TS, Phase ($F(1, 58) = 0.153$, $p = 0.698$), Genotype ($F(1, 18) = 0.0227$, $p = 0.882$), Phase × Genotype ($F(1, 58) = 0.0445$, $p = 0.834$), and all planned comparisons were nonsignificant (all $p_{Holm} = 1$). Bars show means + SEM across sessions. *$p < 0.05$; **$p < 0.01$ (Holm-adjusted).

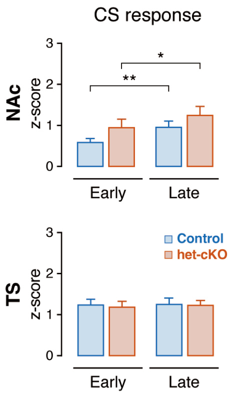
